# A novel Ataxin-3 knock-in mouse model mimics the human SCA3 disease phenotype including neuropathological, behavioral, and transcriptional abnormalities

**DOI:** 10.1101/2020.02.28.968024

**Authors:** Eva Haas, Rana D. Incebacak, Thomas Hentrich, Yacine Maringer, Thorsten Schmidt, Frank Zimmermann, Nicolas Casadei, James D. Mills, Eleonora Aronica, Olaf Riess, Julia M. Schulze-Hentrich, Jeannette Hübener-Schmid

## Abstract

**Background:** Spinocerebellar ataxia type 3 is the most common autosomal dominant inherited ataxia worldwide and is caused by a CAG repeat expansion in the *Ataxin-3* gene resulting in a polyQ expansion in the corresponding protein. The disease is characterized by neuropathological (aggregate formation, cell loss), phenotypical (gait instability, body weight reduction), and specific transcriptional changes in affected brain regions. So far, there is no mouse model available representing all the different aspects of the disease, yet highly needed to gain a better understanding of the disease pathomechanism.

**Methods:** Here, we characterized a novel Ataxin-3 knock-in mouse model, expressing either a heterozygous or homozygous expansion of 304 CAG/CAAs in the murine *Ataxin-3* locus using biochemical, behavioral, and transcriptomic approaches. Further, we compared the transcriptional changes of the knock-in mice to those found in human SCA3 patients, to evaluate the comparability of our model.

**Results:** The novel Ataxin-3 knock-in mouse is characterized by the expression of a polyQ-expansion in the murine Ataxin-3 protein, leading to massive aggregate formation, especially in brain regions known to be vulnerable in SCA3 patients, and impairment of Purkinje cells. Along these neuropathological changes, mice showed a reduction in body weight accompanied by gait and balance instability. Transcriptomic analysis of cerebellar tissue revealed age-dependent differential expression, enriched for genes attributed to myelinating oligodendrocytes. Comparing these transcriptional changes with those found in cerebellar tissue of SCA3 patients, we discovered an overlap of differentially expressed genes pointing towards similar gene expression perturbances in several genes linked to myelin sheaths and myelinating oligodendrocytes.

**Conclusion:** The novel Ataxin-3 knock-in model shares neuropathological, behavioral, and transcriptomic features with human SCA3 patients and, therefore, represents an ideal model to investigate early-onset developments, therapy studies, or longitudinal biomarker alterations.

## Background

Spinocerebellar ataxia type 3 (SCA3) or Machado-Joseph disease (MJD) is the most common inherited ataxia worldwide. It is caused by a CAG repeat expansion in the *ATAXIN-3 (ATXN3)* gene in exon 10, leading to an expanded polyglutamine (polyQ) stretch in the corresponding protein, the deubiquitinase ATXN3 (Kawaguchi *et al*. 1994, Burnett *et al*. 2003). Therefore, it is one of nine so-called polyglutamine (polyQ) diseases (SCA1, 2, 3, 6, 7 and 17, Huntington disease (HD), spinobulbar muscular atrophy, and dentatorubral pallidoluysian atrophy) (Adegbuyiro *et al*. 2017). While healthy individuals present with 13-41 CAG repeats, SCA3 patients show an expansion of 55-86 CAGs in one allele of the *ATXN3* gene (Riess *et al*. 2008). The resulting glutamine expansion in the ATXN3 protein leads to ubiquitin-positive neuronal inclusions, also containing the disease protein ATXN3, demonstrating a typical hallmark of SCA3. While these inclusions may not be toxic (Arrasate *et al*. 2004, Kurosawa *et al*. 2015), misfolding and aggregation of the disease protein is proposed to be the central cause of the disease pathogenesis in polyQ diseases (Hipp *et al*. 2014, Sweeney *et al*. 2017). The age-dependent intraneural accumulation and aggregation of the expanded protein is associated with neuronal dysfunction and cell loss, predominantly of the brainstem, cerebellum, and spinal cord (Schmidt *et al*. 1998, Yamada *et al*. 2004). In most cases, the symptomatic phase of the disease occurs in the third or fourth decade of life and is characterized by motor abnormalities including gait ataxia, ocular symptoms, and cognitive disturbances later in life (Riess *et al*. 2008). So far, there are neither preventive treatments nor good early-onset nor progression biomarkers available (Saute *et al*. 2012).

Mouse models are important tools to investigate disease pathomechanisms, to discover treatment possibilities, or to study longitudinal disease progression and biomarker changes. While several SCA3 mouse models are available, the majority of these have the disadvantage of expressing cDNA under the control of artificial promoters, e.g. CMV, L7, rHtt, PrP or Purkinje cell (PC) specific promotors (Ikeda *et al*. 1996, Goti *et al*. 2004, Bichelmeier *et al*. 2007, Boy *et al*. 2009, Boy *et al*. 2010, Silva-Fernandes *et al*. 2010). Further, a genetrap model, expressing only the N-terminal part of the murine Atxn3 protein, not containing a CAG repeat, presented with an ataxia-like phenotype (Hubener *et al*. 2011). Although these models enabled the investigation of selected SCA3 features, they have certain disadvantages like containing excessive numbers of transgene copies, unnatural expression patterns, incomplete protein expression, the lack of regulatory sequences, or even a combination of these drawbacks. In 2002, Cemal and colleagues presented the first full-gene model expressing the human *ATXN3* gene with all regulatory sequences, yet still expressing the endogenous mouse *Atxn3* gene (Cemal *et al*. 2002). In 2015, the first Atxn3 knock-in (KI) models for SCA3 were introduced. These models hold the advantage of expressing a CAG expansion in the murine *Atxn*3 locus under endogenous regulatory elements and at physiological levels. Ramani and colleagues presented a KI line containing 82 CAG repeats in the murine *Atxn3* locus (Ramani *et al*. 2015, Ramani *et al*. 2017), while Switonski and colleagues generated a mouse in which the murine *Atxn3* gene was replaced by a humanized version with 91 CAG repeats (Switonski *et al*. 2015). Although these mice displayed aggregate formation and a mild phenotype, they did not reflect the complete human SCA3 phenotype. We know from animal models of HD, that it is not sufficient to introduce the same CAG expansion length in mice or rats that is found in the pathogenic human *HTT* gene. Instead several folds of this length are required, to trigger a phenotype (Farshim *et al*. 2018). Thus, we concluded that it is necessary to introduce a much longer CAG repeat in murine *Atxn3* than observed in human patients, to provoke a phenotype.

Therefore, we introduce a novel SCA3 KI mouse model expressing a hyper-expansion of 304 CAACAGs either in one or in both alleles. This new line is characterized by a massive formation of ubiquitin (Ub)-positive Atxn3 aggregates in brain regions vulnerable in SCA3 patients, accompanied by a mild loss of Purkinje cells. Further, mice showed a reduction in body weight, size, and additional physical changes accompanied by a motor phenotype including gait and balance instabilities as well as foot print abnormalities comparable to human SCA3 clinical features. Moreover, cerebellar transcriptome analysis revealed an age-dependent increase in differentially expressed genes (DEGs) clustering in distinct gene expression patterns. When comparing these changes in mice to *post-mortem* cerebellar data from SCA3 patients, an intriguing overlap of DEGs associated with myelinating oligodendrocytes became apparent, indicating shared perturbations between both organisms.

Taken all these findings together, we present a new Atxn3 KI mouse model mimicing the human patient SCA3 phenotype on a behavioral, neuropathological, and transcriptomic level, enabling the analysis of early onset events, longitudinal developments, as well as treatment and biomarker progression.

## Material and Methods

### Ethical use of animals

All mice were maintained by animal care staff and veterinarians of the University of Tuebingen. All procedures were performed according to the German Animal Welfare Act and the guidelines of the Federation of European Laboratory Animal Science Associations, based on European Union legislation (Directive 2010/63/EU). Animal experiments were approved by the local ethics committee (Regierungspraesidium Tuebingen (HG3/13)).

### Ethical use of human tissue

All the work involving human tissue has been carried out in accordance with the Code of Ethics of the World Medical Association (Declaration of Helsinki) and with national legislation as well as our institutional guidelines.

### Generation of KI lines

The KI founder line was generated by Zink-finger technology (Zn-finger) as described previously (Carbery *et al*. 2010). The DNA binding domains of the Zn-finger nucleases were located 3’ and 5’of the CAA(CAG)_5_ region (binding sequence: 5’-acagatattcacgtttgaatgtttcaggCAACAGCAGCAGCAGCAG-gaggtagaccgacctggacccctttcat-3’) in the murine *Atxn3* gene. The associated restriction site generated a double–strand-break (DSB) within the sequence. A donor vector with (CAACAGCAG)_48_ repeats, flanked by 800 bp up- and down-stream of the C57Bl/6 CAA(CAG)_5_ region was used to insert the specific mutation by homologous recombination (HR).

### Genotyping by PCR

DNA was extracted from mouse ear biopsies using High Pure PCR Template Preparation Kit (Roche, Mannheim, Germany) according to the manufacturer’s instructions. Extracted DNA was used for genotyping by polymerase chain reaction (PCR) (Taq DNA polymerase, Qiagen, Hilden, Germay), using primers flanking the endogenous CAG repeat in exon 10 of the murine *Atxn3* gene (primer sequences, Table 1) and the following PCR conditions: 1 cycle 95° C/5 min; 35 cycles of 95° C/30 sec, 55° C/30 sec, 72° C/60 sec; 1 cycle 72° C/10 min. PCR products were separated on a 1% agarose gel and visualized using a 1% ethidium bromide solution (Carl Roth, Karlsruhe, Germany).

**Table 1:**
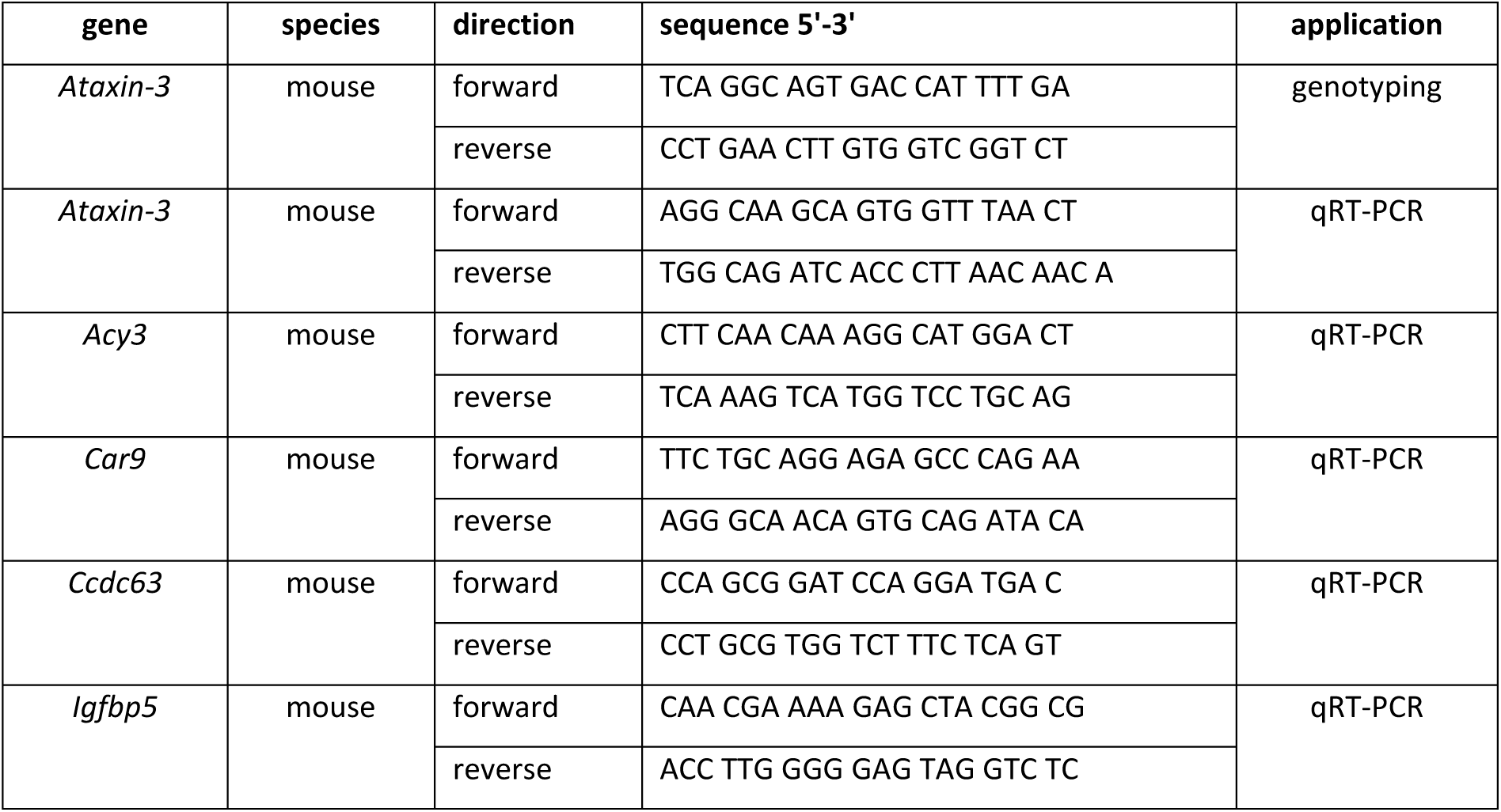
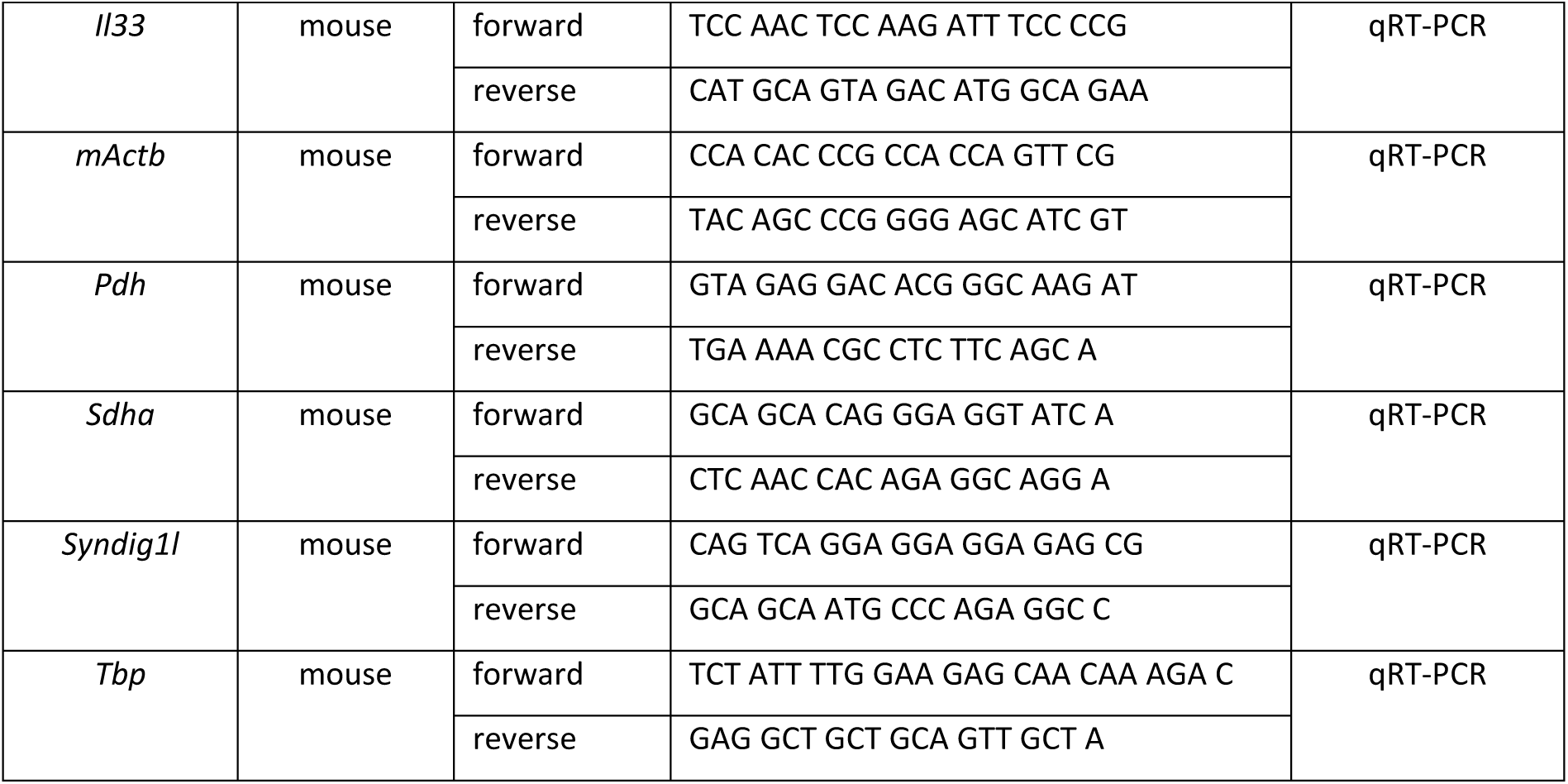
Primer sequences used for genotyping and qRT-PCRs.

### Sequencing of CAG expansion by Sanger Sequencing and determination of CAG length by fragment length analysis

To analyze the right order of the CAACAGCAG expansion Sanger Sequencing was performed. After amplifying the CAG region by PCR (genotyping conditions), the PCR product was purified using the Qiaquick PCR Purification Kit (Qiagen, Hilden, Germany) according to the manufacturer’s instructions. Sequencing was performed with the BigDye Terminator v3.1 Cycle Sequencing Kit (Applied Biosystems, Waltham, USA) using 15-30 ng purified PCR product and the following PCR conditions: 1 cycle 96° C/2 min; 30 cycles of 96° C/10 sec, 55° C/5 sec, 60° C/3 min; 1 cycle 60° C/10 min. Sequencing was performed with either forward or reverse primer, separately. Sequencing products were cleaned up using magnetic beads (CDTR-0050; CleanNA, Waddinxveen, The Netherlands) and according to the Agencourt AMpure PCR purification (Beckman Coulter, Brea, USA) protocol. Separation of the sequences was performed on the ABI 3730xl DNA Analyzer (injection time: 5 sec, run duration: 40 min, separation gel: POP7, Applied Biosystems, Waltham, USA).

Capillary electrophoresis was used to determine the fragment length of the PCR products. A total of 25 μl of the PCR product was cleaned using 1X Agencourt AMPure XP beads (Beckman Coulter, Brea, USA) and eluted in 25 μl TE buffer. Concentration was measured using Qubit dsDNA HS (Cat No Q32854, ThermoFisher Scientific, Waltham, USA). Fragment size was determined by loading 1 ng/μl cleaned up PCR product on the Fragment Analyzer System (Agilent Technologies, Santa Clara, USA) using the qualitative DNA Kit dsDNA Reagent 35-5000bp (Cat No. DNF-915, Agilent Technologies, Santa Clara, USA).

### Animal housing and general health assessment

All animals were housed under standard conditions in type II long cages. After weaning, they were kept with a maximum of five animals of the same gender per cage, avoiding animals housing alone and without enrichment. They were maintained within a 12 hour light-dark cycle and had access to food and water *ad libitum*. Measurement of body weight and a thorough inspection of the health status of the animals were performed every alternate week.

### Accelerating RotaRod test

Twelve to fifteen animals per genotype (WT/WT or KI mice WT/304Q and 304Q/304Q) were placed on an accelerating RotaRod (TSE-Systems, Bad Homburg, Germany) on four consecutive days. Day one to three were training sessions, each with an acceleration from 4 rpm to 16 rpm in 2 min. Two rounds of training were performed per training session. Day four was the test session with two test trials, each test trial with an acceleration from 4 rpm to 40 rpm in 5 min. The time the mice spend on the accelerating rod was recorded. Each training and test session started at the same time of the day and two trials per day were performed. Mice were allowed to rest for 1 hour between trials. Accelerating RotaRod test was performed at 3, 6, 9, 12, 15 and 18 months of age.

### Gait analysis

Gait analysis was performed with the gait analysis system Catwalk 8.1 (Noldus, Wageningen, The Netherlands). Twelve to fifteen animals per genotype were placed individually in a tunnel on a glass plate where the mice could move voluntarily. Each mouse had to perform five runs with minimal run duration of 0.5 sec and a maximum of 10 sec. The maximum variation in speed was 60%. Runs with rearing behavior or a change of direction were automatically discarded. Evaluations of the runs were performed with the Catwalk X software (Noldus, Wageningen, The Netherlands). Gait analysis was assessed at 6, 12 and 18 months of age.

### Tissue preparation for RNA and protein analysis

To investigate protein and RNA levels, WT/304Q and 304Q/304Q KI mice and their WT/WT littermates were sacrificed by CO_2_ inhalation followed by head decapitation (n = 3 per genotype and experimental setup). Brain regions and peripheral organs for protein analysis were immediately dissected and snap-frozen in liquid nitrogen. Cerebellar samples for RNA analysis were stabilized overnight in RNAlater (Qiagen, Hilden, Germany). All samples were stored at −80° C until further usage.

### Tissue homogenate and lysate for protein analyses

For tissue lysate and homogenate preparation, frozen tissue was mechanically homogenized in TES buffer (4% Tris Base pH 7.5, 0.1 mM EDTA, 100 mM Na_2_Cl) containing protease inhibitor (PI) (cOmplete, EDTA-free Protease Inhibitor Cocktail (Roche, Mannheim, Germany)) with the VDI 12 homogenisator (VWR, Darmstadt, Germany). TNES (TES-buffer + 10% Igepal CA-630) was added in a relation 1:50 and incubated on ice for 30 min. Part of the homogenates was stored at –80° C and later used for filter retardation assay. The rest of the homogenates were centrifuged at 13.200 x g for 25 min at 4° C and the supernatants (lysates) were transferred to a new tube. Glycerol was added to a final concentration of 10%. Protein concentration was measured spectrophotometrically by Bradford Assay (Bio-Rad Laboratories, Feldkirchen, Germany).

### Western blot analysis

Western blotting was performed using standard procedures. In short, 4x LDS sample buffer (1 M Tris Base pH 8.5, 2 mM EDTA, 8% LDS, 40% glycerol, 0.025% phenol red) and 100 mM 1,4-dithiothreitol (DTT) (MerckMillipore, Darmstadt, Germany) was added to 30 μg of protein lysates and heat-denatured at 70° C for 10 min. Adjacent, protein samples were separated by electrophoresis using a 8% Bis-Tris gel and MOPS (50 mM MOPS, 50 mM Tris Base pH 7.3, 3.5 mM SDS, 1mM EDTA) or MES (50 mM MES, 50 mM Tris Base pH 7.3, 0.1% SDS, 1 mM EDTA) electrophoresis buffer. Proteins were transferred on Amersham Protran Premium 0.2 μm nitrocellulose membranes (GE Healthcare, Solingen, Germany) a Bicine/Bis-Tris transfer buffer (25 mM Bicine, 25 mM Bis-Tris pH 7.2, 1 mM EDTA, 15% methanol) and a TE22 Transfer Tank (Serva, Heidelberg, Germany) at 80 V for 2 hours by 4° C was used. Afterward, membranes were blocked in 5% skimmed milk powder (Sigma-Aldrich, Munich, Germany) in TBS (Tris-buffered saline) (1 M Tris, 5 M NaCl) for 1 hour at room temperature and incubated with primary antibodies (rabbit anti-Acy3, 1:500, LS-C401171, LSBio; mouse anti-Ataxin-3, 1:2500, clone 1H9, MAB5360, MerckMillipore, Darmstadt, Germany; rabbit anti-Car9, 1:2000, NB100-417, novusbio, Littleton, USA; rabbit anti-Ccdc63, 1:500, LS-C111466, LSBio, Seattle, USA; rabbit anti-Igfbp5, 1:400, LS-C294665, LSBio, Seattle, USA; rabbit anti-IL-33, 1:400, LS-C294906, LSBio, Seattle, USA) overnight at 4° C. Subsequently, membranes were washed with TBST (TBS + 0.1% Tween 20) and incubated with secondary antibodies (peroxidase affinePure donkey anti-mouse IgG H+L, 1:12000, 715-035-150, Jackson ImmunoResearch, Ely, United Kingdom; IRdye 800CW goat anti-mouse IgG (H+L) 1:1000, 926-32210, LI-COR, Bad Homburg, Germany; IRdye 800CW goat anti-rabbit IgG (H+L) 1:1000, 926-32211, LI-COR, Bad Homburg, Germany; goat anti-rabbit IgG H&L (HRP), 1:10.000, ab97051, Abcam, Cambridge, United Kingdom) for 1.5 hours at room temperature (RT). In case of chemiluminescent detection, WesternBright Chemilumineszenz Substrat Sirius (Biozym, Hessisch Oldendorf, Germany) was used in the ratio 1:1:1 with one part TBS. Fluorescence or chemiluminescent signal detection was performed on the LI-COR ODYSSEY FC and quantified using the ODYSSEY Server software version 4.1 (LI-COR, Bad Homburg, Germany).

### Filter retardation assay

Filter retardation assay was used to detect SDS-insoluble Atxn3 and ubiquitin (Ub) species in brain tissue homogenates. Therefore, 12.5 μg of total protein was diluted in Dulbecco’s phosphate-buffered saline (DPBS) (Gibco, Waltham, USA) with 2% SDS and 50 mM DTT and heat denatured at 95° C for 5 min. To avoid precipitation of SDS cool down was performed at room temperature. After equilibrating Amersham Protran Premium 0.45 μm nitrocellulose membranes (GE Healthcare, Solingen, Germany) with DPBS containing 0.1% SDS samples were filtered through the membranes by using a Minifold® II Slot Blot System (Whatman, Maidstone, United Kingdom). Slots were washed with one volume of DPBS before washing the whole membrane with TBS and blocking with 5% skimmed milk powder (Sigma-Aldrich, Munich, Germany) in TBS for 1 hour. Membranes were incubated overnight at 4° C with primary antibodies (mouse-anti-Ataxin-3, 1:2500, clone 1H9, MAB5360, MerckMillipore, Darmstadt, Germany; rabbit-anti-Ubiquitin, 1:500, Z0458, Dako, Jena, Germany). After washing the membranes were incubated with secondary antibodies (peroxidase affinePure donkey anti-mouse IgG H+L, 1:12000, 715-035-150, Jackson ImmunoResearch, Ely, United Kingdom; IRdye 800CW goat anti-mouse IgG (H+L) 1:1000, 926-32210, LI-COR, Bad Homburg, Germany; IRdye 800CW goat anti-rabbit IgG (H+L) 1:1000, 926-32211, LI-COR, Bad Homburg, Germany; goat anti-rabbit IgG H&L (HRP), 1:10.000, ab97051, Abcam, Cambridge, United Kingdom) for 1.5 hours at room temperature and detected as described in the section “Western blot analysis”.

### Time-Resolved Fluorescence Resonance Energy Transfer (TR-FRET) immunoassay

TR-FRET immunoassay was used to measure total and expanded soluble Atxn3 protein levels in whole brain homogenates of 3- and 18-month-old mice. These immunoassays are based on an energy transfer of an excited donor fluorophore towards an acceptor fluorophore, which is only possible when both bind in close spatial proximity (Degorce *et al*. 2009). Terbium cryptate (tb) was used as donor fluorophore and d2 as acceptor fluorophore. To detect total soluble Atxn3 protein, 10 ng/μl anti-Atxn3-d2 (mouse-anti-Ataxin-3, clone 1H9, MAB5360, MerckMillipore, Darmstadt, Germany) and 0.5 ng/μl anti-Atxn3-(N-terminal)-tb (rabbit-anti-Ataxin-3, ab96316, Abcam, Cambridge, United Kingdom) were mixed in 1x detection buffer (in 50 mm NaH2 PO4, 400 mm NaF, 0.1% BSA, 0.05% Tween-20). To measure the amount of polyQ-expanded Atxn3, 3 ng/μl anti-polyQ-d2 (mouse-anti-Polyglutamine-Expansion Diseases Marker Antibody, clone 5TF1-1C2, MAB1574, MerckMillipore, Darmstadt, Germany) and 1 ng/μl anti-Atxn3-tb (mouse-anti-Ataxin-3, clone 1H9, MAB5360, MerckMillipore) were diluted in detection buffer. For both measurements, 5 μl of whole brain homogenates of 3- and 18-month-old mice were incubated with 1 μl of the respective antibody mix and incubated at 4°C for 22 hours. Donor fluorescence was measured at 615 nm and acceptor fluorescence at 665 nm with EnVision 2105 multimode plate reader unit (PerkinElmer, Waltham, USA). For quantification, the acceptor to donor fluorescence ratio was calculated and corrected to total protein amount and negative control. The resulting fluorescence signal is proportional to the protein concentration.

### Immunohistochemistry and microscopy

To obtain sections, mice were transcardially perfused with cold PBS and 4% paraformaldehyde (PFA) and the brains were post-fixed in 4% PFA overnight at 4° C and subsequently embedded in paraffin. For immunohistochemical staining paraffin-embedded brains were cut in 7 μm thick sagittal sections with the Leica RM2155 microtome (Leica, Wetzlar, Germany)and were rehydrated in xylene and a graded alcohol series using the Leica autostainer XL (Leica, Wetzlar, Germany). Microwave treatment with 10 mM sodium citrate and 10 mM citric acid for 15 min and washing with phosphate buffered saline (PBS) was performed.

For the immunohistochemical staining, the endogenous peroxidase was blocked by using 1.6% peroxidase (Sigma-Aldrich, Munich, Germany). After washing with PBS sections were blocked in 5% normal goat serum (NGS) (Vector Laboratories, Burlingame, USA) in PBS supplemented with 0.3% Triton X-100 (Carl Roth, Karlsruhe, Germany) at RT for 45 min. After washing with PBS, sections were incubated with primary antibody mouse anti-Ataxin-3 (clone 1H9, 1:400, MerckMillipore, Darmstadt, Germany) or rabbit anti-Calbindin (D28-K, 1:1000, Swant, Marly, Switzerland) diluted in PBS containing 15% NGS overnight at 4°C in a humid chamber. After washing with PBS, biotinylated secondary antibody goat anti-mouse (1:200, Vector Laboratories, Burlingame, USA) or goat anti-rabbit (1:200, Vector Laboratories, Burlingame, USA) were incubated on the sections for 1 hour at room temperature. In parallel, the avidin-biotin-complex (ABC) (Vector Laboratories, Burlingame, USA) was prepared and after secondary antibody incubation, ABC was incubated on the sections for two hours at room temperature. After washing with PBS the substrate 3,3′-Diaminobenzidin (DAB, Sigma-Aldrich, Munich, Germany) was added to the sections and the reaction was stopped in distilled water when the desired degree of staining was reached. After dehydrating the sections were mounted with CV Ultra mounting media (Leica, Wetzlar, Germany).

Imaging of the sections occurred with the Axioplan2 imaging microscope using an Axio-Cam HR color digital cam, a 20x Plan Neofluar and 63x Plan/Apochromat objective and the AxioVison SE64 Rel. 4.9 software (all Zeiss, Oberkochen, Germany).

Overview images were taken with the AxioScan.Z1 in 40-fold magnification and analyzed with the Zen2011 software (all Zeiss, Oberkochen, Germany).

### RNA Sequencing of human and mouse cerebellar RNA

For mice, total RNA, microRNA, and DNA were extracted simultaneously using the AllPrep DNA/RNA/microRNA Universal Kit (Qiagen, Hilden, Germany) by using the manufacturer’s protocol. RNA was isolated from cerebellar tissue (n = 5 WT/WT and 304Q/304Q mice) at a pre-symptomatic (2 months of age) and symptomatic (12 months of age) time point. Samples with very high RNA integrity numbers (RIN > 8) were selected for library construction. A total of 100 ng of total RNA was subjected to polyA enrichment and cDNA libraries were constructed using the resulting mRNA and the TruSeq Stranded mRNA (Cat No 20020595, Illumina, San Diego, USA). Libraries were sequenced as paired-end 101 bp reads on a NovaSeq6000 (Illumina, San Diego, USA) with a depth of 16 - 38 million reads each. Library preparation and sequencing procedures were performed by the same individual and a design aimed to minimize technical batch effects was chosen.

For human samples, frozen tissue was homogenized in Qiazol Lysis Reagent (Qiagen Benelux, Venlo, The Netherlands). The total RNA including the miRNA fraction was isolated using a miR-Neasy Mini kit (Qiagen Benelux, Venlo, the Netherlands) according to manufacturer’s instructions. RNA was isolated from six male SCA3 patients (mean age: 66.8 ± 7.7 years) and six male controls not affected by SCA3 (mean age: 64.3 ± 16.3 years). The human samples presented with a lower quality RNA integrity number (RIN > 4 and RIN < 7, DV200 >60 %) as well as a low quantity of RNA. Library preparation was performed by capture of sequence-specific coding RNA using 40 ng of total RNA and the TruSeq RNA Access Library Prep Kit (Cat No RS-301-2001, Illumina, San Diego, USA). Libraries were sequenced as paired-end 68 bp reads on a HiSeq2000 (Illumina, San Diego, USA) with a depth of approximately 25 - 100 million reads each.

### Analysis of RNA Sequencing

Quality of the RNA sequencing (RNA-seq) data was assessed using *FastQC* (v0.11.4) (http://www.bioinformatics.babraham.ac.uk/projects/fastqc) to identify sequencing cycles with low average quality, adaptor contamination, or repetitive sequences from PCR amplification before aligning reads with *STAR* (v2.5.4b) (Dobin *et al*. 2013) against the *Ensembl* M. musculus and H. sapiens genome v91 allowing gapped alignments to account for splicing. Alignment quality was analyzed using *samtools* (v1.1) (Li *et al*. 2009). Normalized read counts for all genes were obtained using *Rsubread* (v2.0.0) and *DESeq2* (v1.18.1) (Love *et al*. 2014). Transcripts covered with less than 50 reads were excluded from subsequent analyses leaving 12.108 (mouse) and 14.893 (human) genes for determining differential expression. The factorial design of the experiment was captured in a general linearized model defining gene expression as a function of genotype, age, and interaction of both. Significance thresholds were set to | log_2_ FC | ≥ 0.5 and BH-adjusted *p*-value ≤ 0.1. Surrogate variable analysis (*sva*, v3.26.0) (Leek *et al*. 2012) was used to minimize unwanted variation between samples. Gene-level abundances were derived from *DESeq2* as normalized read counts and used for calculating the log2-transformed expression changes of the expression heatmap and centroids. Ratios were relative to mean expression in WT/WT_2m_. The *sizeFactor*-normalized counts provided by *DESeq2* also went into calculating nRPKMs (normalized Reads Per Kilobase per Million total reads) as a measure of relative gene expression as motivated before (Srinivasan *et al*. 2016). Orthologous genes between mouse and human were determined with the *biomaRt* package on the v91 of the *Ensembl* genome annotations. Cell type-specific expression data for mouse and human were adapted from Zhang *et al*. (2014) and Kuhn *et al*. (2012), respectively. *WebGestalt* was employed to identify overrepresented molecular functions among *Gene Ontology* terms (Wang *et al*. 2013). KI raw sequencing files are available through GEO under accession number: GSE145613. Human RNA-Seq data set has been deposited at the European Genome-phenome Archive (EGA), which is hosted by the EBI and the CRG, under the accession number: EGAS00001004241.

### Quantitative Reverse Transcription PCR (qRT-PCR)

qRT-PCR was performed for validation of the differentially expressed genes *Acy3, Atxn3, Car9, Ccdc63, Igfbp5, Il33*, and *Syndig1l*. 500 ng purified RNA was transcribed into cDNA using QuantiTect Reverse Transcription Kit (Qiagen, Hilden, Germany). For each gene of interest, 2 μl diluted cDNA (1:20) were added to SYBR Green PCR Master Mix (Qiagen, Hilden, Germany) and 10 μM of both forward and reverse primer (Primer Sequences Table 1). qRT-PCR was run on the LightCycler 480 II (Roche, Mannheim, Germany). The relative gene expression was calculated by normalization to housekeeping genes *SDHA, PDH, mACTB* and *TBP* (Primer Sequences Table 1).

### Statistical analysis

For statistical analysis, GraphPad Prism 6.0 for Windows (GraphPad Software Inc., San Diego, USA) was used. Significance of datasets was determined using two-tailed Student’s t-test comparing either WT/WT animals with each KI group (significances marked by *) or the mice of the KI lines WT/304Q and 304Q/304Q (marked by #) respectively. For behavior analysis Welsh Correction was applied. P values with less than 0.05 were considered statistically significant with */# p < 0.05, **/## p < 0.01 and ***/### p < 0.001. All values are shown as mean ± standard derivation, SEM.

Enrichments for cell types are based on Fisher’s exact tests (R *stats* package v3.6.2). Shifts in cell-type composition were assessed using Mann-Whitney U tests (R *stats* package v3.6.2).

## Results

### Zinc-finger generated Ataxin-3 KI line expressed expanded Atxn3

Our goal was the generation of a new SCA3 KI mouse line with a hyper-expanded polyQ stretch in the murine *Atxn3* gene under control of all endogens regulatory elements to trigger a neurological phenotype comparable to human SCA3 patients. Thus, we used Zinc-finger (Zn-finger) technology (Carbery *et al*. 2010) to introduce the CAG expansion into the murine *Atxn3* locus in mice with C57BL/6 background (Figure 1A). Here, the DNA binding domains I and II recognized sequences flanking the endogenous murine sequence CAA(CAG)_5_ and introduced a double strand break (DSB). For homologous recombination (HR), we used a donor vector including the CAACAGCAG expansion with the interrupted sequence (CAACAGCAG)_48_ flanked with 800 bp of the up- and downstream region of the *Atxn3*. Due to the repetitive sequence in the donor vector, we obtained a founder line containing 304 CAACAG repeats, leading to the corresponding length of polyQ expansion in the Atxn3 protein (in the following called WT/304Q mice) (Figure 1A). To analyze a potential gene dosage effect, homozygous mice (304Q/304Q) were bred as well. Both genotypes were viable, fertile, and showed no signs of reduced survival. Protein analysis confirmed the expression of non-expanded (WT) and expanded Atxn3 in whole brain lysates for the heterozygous line at an early (3 months of age) and late (18 months of age) time point (Additional file 1: Figure S1A + S1B). Non-expanded Atxn3 was detected with a molecular weight of 42 kDa in WT/WT and WT/304Q mice, and expanded Atxn3 was found to be detected with a molecular weight of about 180 kDa in WT/304Q and 304Q/304Q mice due to the increased polyQ size in the Atxn3 protein. Analysis of whole brain lysates of homozygous 304Q/304Q mice showed expression of expanded Atxn3 and complete loss of non-expanded Atxn3 (Additional file 1: Figure S1A + S1B).

**Figure 1:**
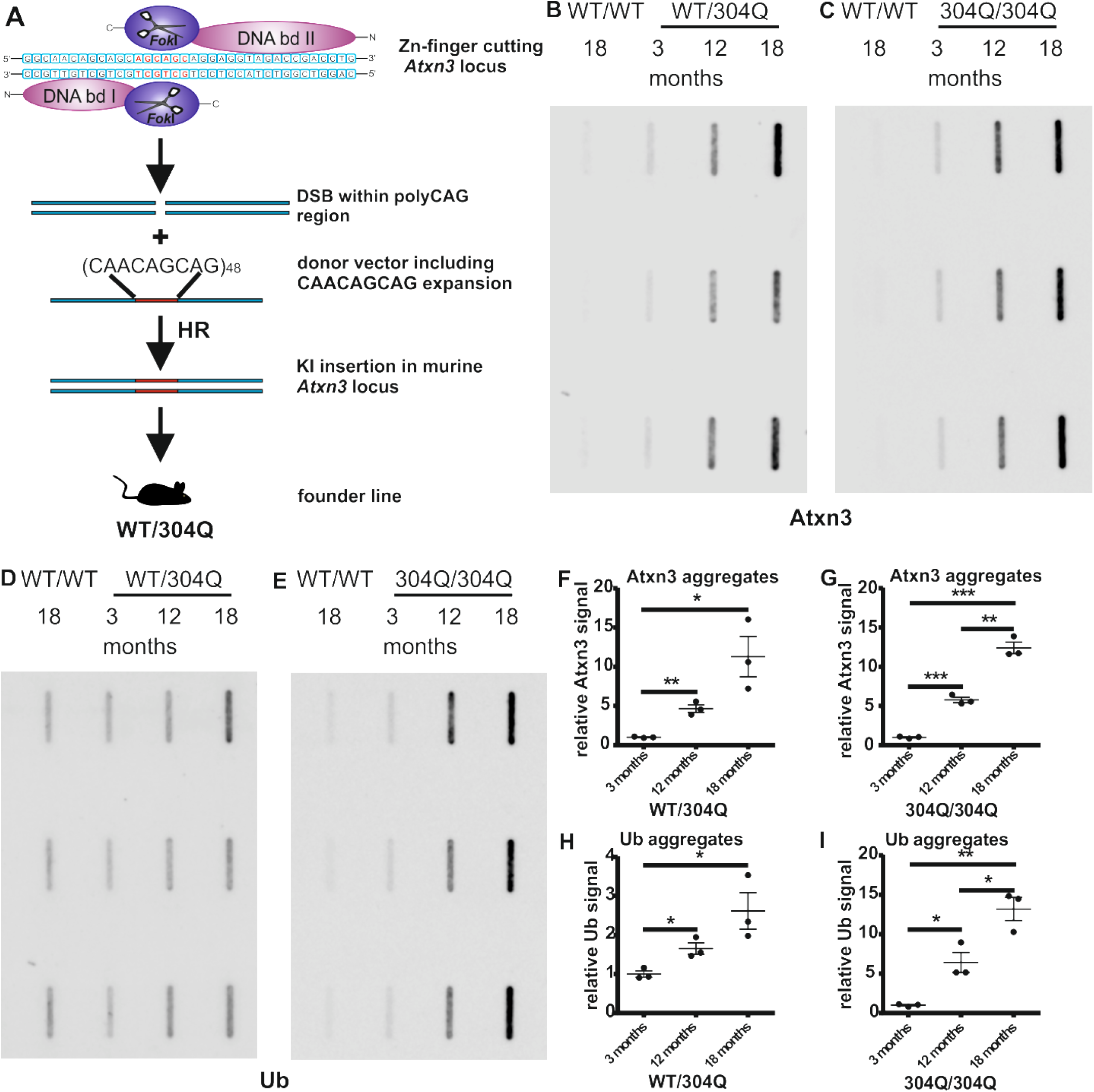
Zinc-finger generated Atxn3 KI mice showed Atxn3 and ubiquitin-positive aggregates. (A) Schematic representation of KI generation. Zinc-finger nuclease DNA binding domains recognize sequences in the mouse genome to introduce a double-strand break (DSB) in the mouse *Atxn3* CAG repeat region. Using a donor vector with (CAACAGCAG)_48_ as template for homologous recombination (HR) a mouse line containing 304 CAG repeats in the *Atxn3* locus was generated. In WT/304Q (B+F) and 304Q/304Q (C+G) mice, a significant increase in the amount of Atxn3 positive insoluble aggregates was observed over time. In WT/304Q (D+H) and 304Q/304Q (E+I) aggregates were also stained positive for ubiquitin, and ubiquitin signal was increasing over time. n = 3, all signal values normalized to signal of 3-month-old WT/304Q or 304Q/304Q mice, respectively. Two-tailed Student’s t-test, * p < 0.05, ** p < 0.01, *** p < 0.001.

Further, protein analysis confirmed the expression of expanded Atxn3 in a wide range of tissues such as heart, lung, liver (Additional file 1: Figure S1C), and spleen (Additional file 1: Figure S1D). No expression in kidney and muscle tissue was observed (Additional file 1: Figure S1D).

### Accumulation of ubiquitin-positive Atxn3 aggregates in Atxn3 KI mice with 304 polyQ expansion

One hallmark of SCA3 is the formation of Atxn3- and ubiquitin (Ub)-positive protein aggregates (Schmidt et al., 2002). To demonstrate this feature in our KI mice, we investigated the formation of detergent-insoluble aggregates in whole brain homogenates by filter retardation assay. For KI mice expressing expanded Atxn3, a massive increase of Atxn3-positive aggregates was detectable within 18 months (Figure 1B + 1C, 1F + 1G). Specifically, we observed an increase of 4.6-fold in the first 12 months and a final increase of 11.3-fold at 18 months of age for the amount of Atxn3-positive aggregates in WT/304Q mice compared to 3 months (Figure 1B + 1F). In 304Q/304Q, an increase of 5.8-fold of Atxn3-positive aggregates was observed within 12 months, and an increase of 12.4-fold at 18 months of age (Figure 1C + 1G). In both cases, the formed aggregates were Ub-positive (Figure 1D + 1E), showing mild ubiquitination already with 3 months of age that kept increasing over time. In fact, for the heterozygous WT/304Q mice, we observed an increase of 1.7-fold in the first 12 months and an increase of 2.6-fold at 18 months of age (Figure 1H). This effect was drastically stronger in homozygous 304Q/304Q, where the increase of Ub-positive aggregates was 6.4-fold within 12 months and 13.2-fold at 18 months of age (Figure 1I), respectively.

### Aggregate formation accrued in SCA3 associated brain regions accompanied by Purkinje cell loss

As we saw in the filter retardation assays that there was a massive formation of detergent-insoluble aggregates, we tested which mouse brain areas were most severely affected. Immunohistochemical staining (IHC) for Atxn3 in 3-month-old animals already showed the formation of aggregates in the hippocampus of WT/304Q mice, and in loop III of the cerebellum, the deep cerebellar nuclei (DCN), the pons, and the hippocampus in 304Q/304Q mice (Additional file 2: Figure S2A - S2D). All these areas have been reported to be vulnerable brain regions in SCA3 patients (Schmidt *et al*. 1998, Yamada *et al*. 2004). With 18 months of age, we observed aggregates in all brain areas listed above in both WT/304Q and 304Q/304Q mice (Figure 2A - 2D). While the number of aggregates stayed at a moderate level in the cerebellar loop III (Figure 2A), high numbers of aggregates were detected in the DCN, pons, and hippocampus (Figure 2B - 2D). The massive increase in aggregates in certain areas (e.g. hippocampus) was even visible in sagittal overview images of brains in 18-month-old 304Q/304Q KI mice (Additional file 3: Figure S3). Further, we investigated the state of the cerebellar Purkinje Cells (PCs), a cell type affected in SCA3 patients (Scherzed *et al*. 2012). Staining with Calbindin, a marker for PCs, showed a reduced number of PCs in cerebellar loop III already in 3-month-old 304Q/304Q mice. With 18 months of age, the PCs were shrunken in size in both WT/WT and 304Q/304Q mice compared to 3 months, and again the number of PCs in the KI mice was reduced compared to WT/WT littermates (Figure 2E). A general reduction or shrinkage of cells, measured by layer thickness of granular (GL) and molecular layer (ML) in cerebellar loop IV/V, was not observed in the KI mice (Figure 2F).

**Figure 2:**
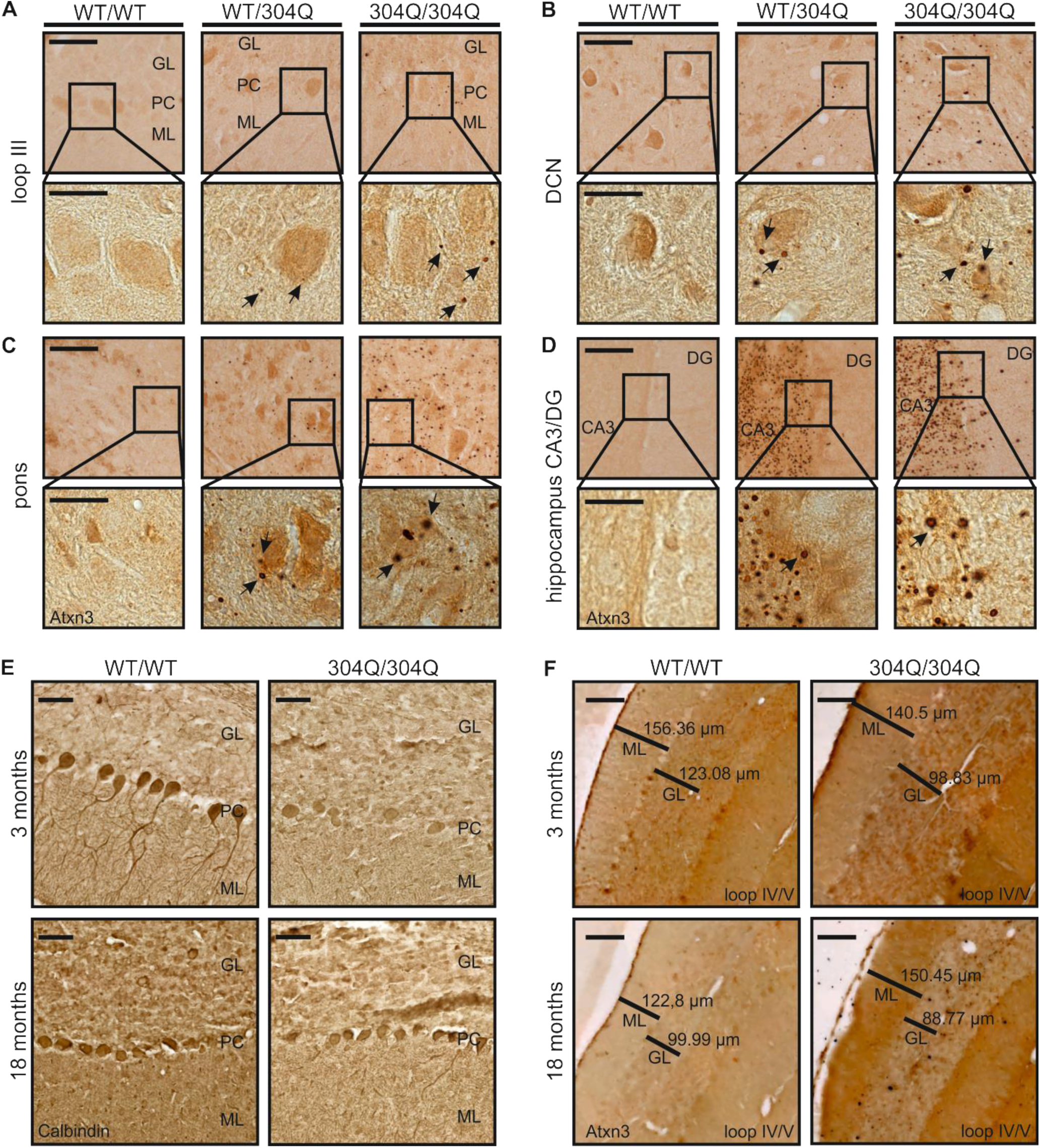
Aggregation of mutant Atxn3 in 18-month-old SCA3 KI mouse brain. Immunohistochemical (IHC) staining using Atxn3-specific antibody (clone 1H9) showed aggregate formation in WT/304Q and 304Q/304Q mice. (A-D) At 18 months, increased diffuse nuclear staining and massive formation of aggregates were detectable in both WT/304Q and 304Q/304Q mice. Most aggregates were found in the DCN (B), the pons (C), and the hippocampal CA3 region (D). (E) Calbindin staining revealed reduced numbers of PCs in 304Q/304Q mice compared to WT/WT littermates in 3- and 18-month-old mice. (F) In cerebellar loop IV/V no genotype- or age-dependent differences in the cerebellar layer thickness were detected (A-E) Scale bar = 50 μm, inset scale bar = 20 μm. (F) Scale bar = 100 μm, Atxn3 positive aggregates are indicated by arrows. (A-D) n=3 (E-F) n = 1, GL = granular layer, PC = Purkinje cells, ML = molecular layer; DG = dentate gyrus; DCN = deep cerebellar nuclei.

### Impaired physical condition and body weight reduction in male mice

As the body mass index (BMI) is known to be reduced in SCA3 patients compared to controls (Saute *et al*. 2011) and to reversely correlate with the length of the expanded polyQ (Saute *et al*. 2012), we investigated whether our SCA3 KI mice display differences in body weight compared to their WT/WT littermates. Therefore, body weight of the mice was measured every other week. For both WT/304Q and 304Q/304Q mice, separating cohorts by gender revealed that male mice (Figure 3A) were affected earlier and more severely by body weight reduction than female mice (Figure 3B). 304Q/304Q males showed significant weight reduction compared to their heterozygous WT/304Q littermates already at 18 weeks of age and to their WT/WT littermates at 20 weeks of age. Heterozygous WT/304Q male mice had a significant reduction in body weight compared to WT/WT animals starting at 48 weeks of age. At the last measurement in our study (at 18 months of age), WT/304Q mice and 304Q/304Q mice had an approximate weight reduction of 24% and 36%, respectively, when compared to WT/WT male mice. In females, we observed a significant reduction in body weight only in 304Q/304Q mice compared to WT/WT littermates starting at 40 weeks of age and compared to their WT/304Q littermates at 42 weeks of age (Figure 3B), but these differences did not reach significance after 58 weeks of age. At the last time point of investigation, the 304Q/304Q mice had an approximate 14% reduction in their body weight compared to WT/WT littermates. Moreover, a reduction in body weight did not represent the only physical change in our newly generated KI mouse lines. Differences in body size (Figure 3C) and posturing (Figure 3D) became apparent for WT/304Q and 304Q/304Q mice. In both, we observed a reduced body size and enhanced hunchback development. Both findings were more pronounced in males (Figure 3C + 3D) and less or not at all detectable in female mice (data not shown).

**Figure 3:**
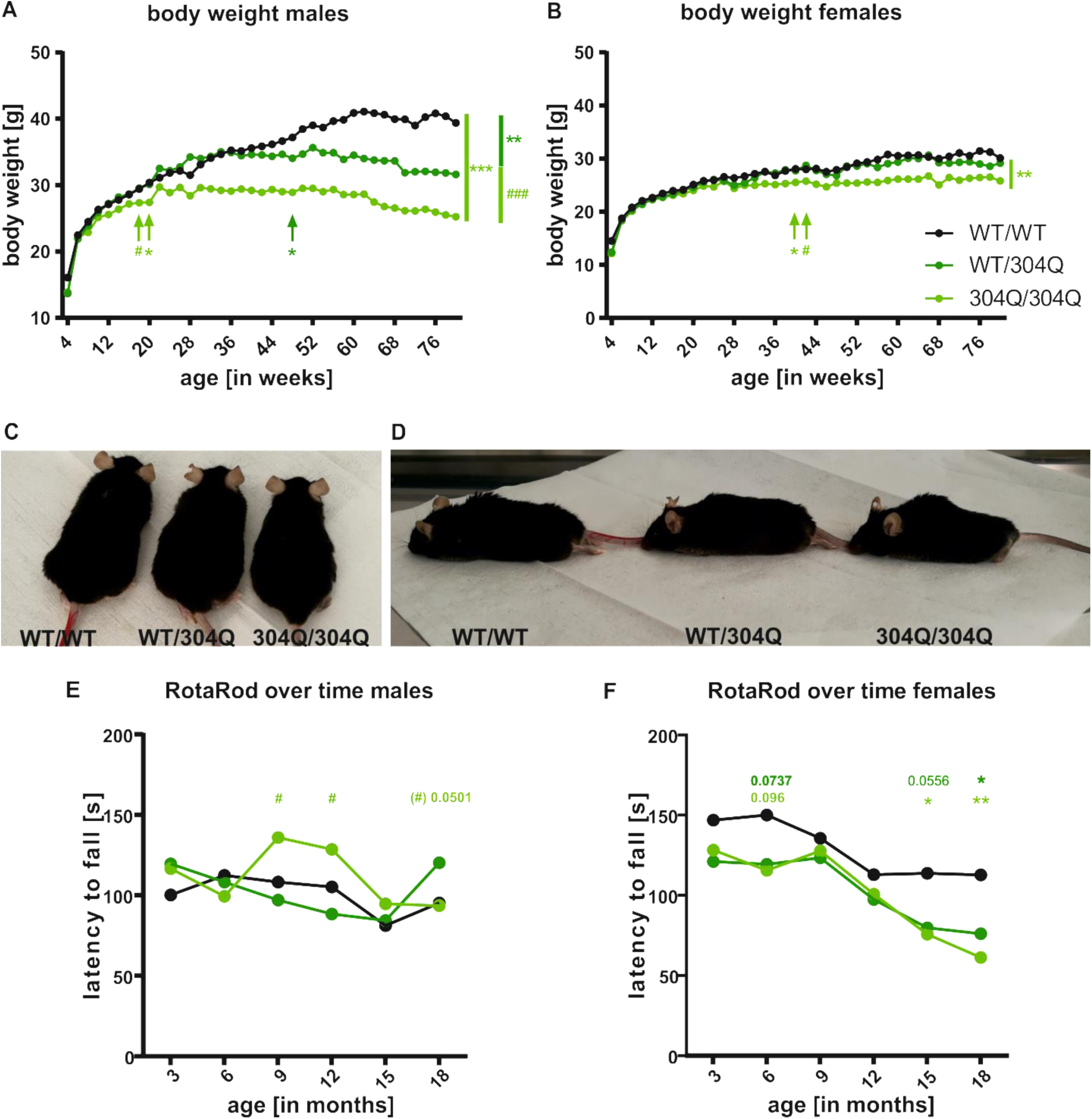
KI mice showed alterations in size, posturing, and body weight accompanied by balance and gait stability. (A) Body weight of male WT/304Q and 304Q/304Q mice is significantly reduced compared to controls after 18 months. (B) In female mice only 304Q/304Q mice showed a reduction in body weight. Beginning of significant differences are indicated by arrows. (C+D) Alterations in size (C) and posturing (D) were well recognizable in 18-month-old male KI mice. (E) 304Q/304Q male mice showed better coordination compared to WT/WT littermates at 9 and 12 months of age. WT/304Q males performed better at 18 months of age than their 304Q/304Q littermates. (F) WT/304Q and 304Q/304Q female mice tend to perform worse than their WT/WT littermates at all measured time points, reaching significance with 15 and 18 months of age. n = 6-8 mice per genotype and gender, two-tailed Student’s t-test with Welsh correction * or # p < 0.05, *** or ### p < 0.001, * comparison WT/WT to KI lines, # comparison WT/304Q to 304Q/304Q, black = WT/WT, dark green = WT/304Q, bright green = 304Q/304Q.

### Female mice present with impaired coordination and stability phenotype while gait analysis was changed in both genders

Changes in body weight also have an effect on the coordination and balancing phenotype of the KI mice. RotaRod analysis conducted to address these features revealed different results for male and female mice. In male mice, no differences between WT/WT and WT/304Q mice were observed, while the 304Q/304Q mice performed better at intermediate time points (9 and 12 months of age) and became comparable to WT/WT and WT/304Q mice in later time points (15 and 18 months) (Figure 3E). In female mice (Figure 3F), WT/304Q and 304Q/304Q showed a tendency for coordination deficits already in young mice (3-month-old) and became significant at later time points (15- and 18-month-old). A detailed presentation of RotaRod analysis for males and females at baseline (3 months) and last measured time point (18 months) demonstrated no coordination differences in young mice (3 months) in neither male (Additional file 4: Figure S4A) nor females (Additional file 4: Figure S4B). In old male mice (Additional file 4: Figure S4C), we observed a tendency for coordination differences in 304Q/304Q mice compared to the WT/304Q mice, while in old female mice (Additional file 4: Figure S4D) we found significantly impaired coordination in WT/304Q and 304Q/304Q mice compared to their WT/WT littermates.

Gait analysis, which seemed not to be as strongly affected by the body weight, showed significant differences in several gait parameters determined by the Catwalk (Noldus) gait analysis system in 18-month-old mice for the pooled cohort without gender separation. For step cycle (SC) analysis, measuring the time it takes a mouse to remove the paw from the surface and to put it back, we observed for the right front paw (Figure 4A) an increased SC in 304Q/304Q mice compared to their WT/WT and WT/304Q littermates. For the right hind paw (Figure 4B) both lines, WT/304Q and 304Q/304Q, showed a significantly increased SC time compared to WT/WT mice. The print area (PA) describes the area of the paw that touches the surface during walking. For the right front paw (Figure 4C) we observed reduced PA in 304Q/304Q mice compared to WT/WT mice and for the right hind paw (Figure 4D), the PA of 304Q/304Q mice were significantly reduced to WT/WT and WT/304Q mice, respectively. Additionally, the base of support (BOS) of the hind paws (Figure 4E), describing the distance between the left and right hind paw and giving also an indication of the balancing capabilities of the mice, was investigated. Here, we observed a significant increase in the BOS for WT/304Q and 304Q/304Q mice, indicating that they need a wider stand for stabilization than their WT/WT littermates. Comparable results were obtained for the left paw (data not shown). Figure 4F shows representative images of the Catwalk quantification. The reduced PA for the 304Q/304Q mice and the decreased BOS for WT/304Q and 304Q/304Q can be seen.

**Figure 4:**
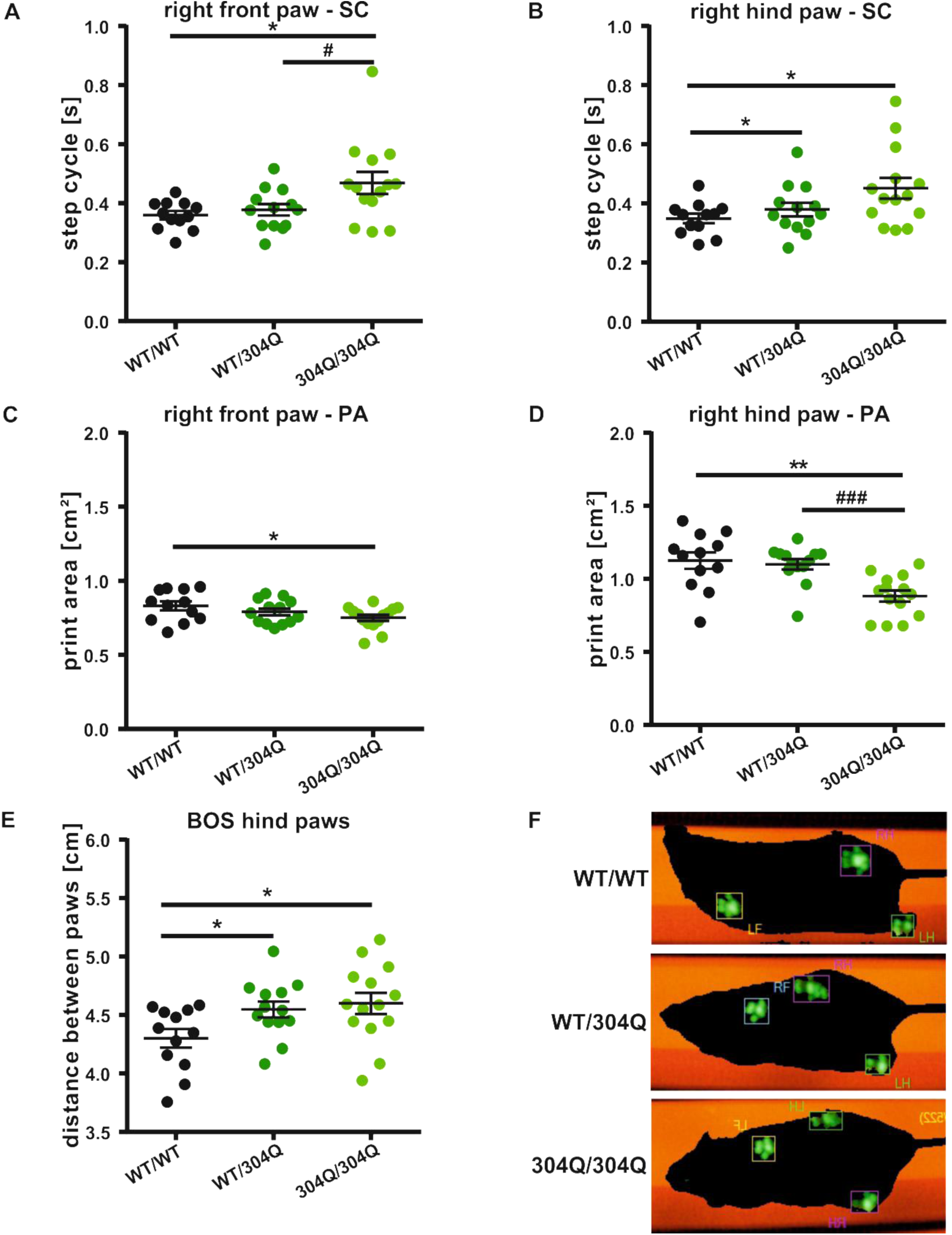
Altered paw positioning in 18-month-old WT/304Q and 304Q/304Q KI mice. (A-B) Step cycle (SC) was increased for 304Q/304Q and WT/304Q mice in the right front paw (A) for and in the hind paw (B) for 304Q/304Q mice compared to WT/WT and WT/304Q littermates. (C-D) Print area (PA) of the right front (C), and hind paw (D) was significantly reduced in 304Q/304Q mice. For the hind paw, this reduction was also significantly different to WT/304Q males. (E) Base of support (BOS) was significantly increased in KI mice compared to WT/WT mice. (F) Visualization of the foot print analysis showed an increase in BOS for WT/304Q and 304Q/304Q mice. n = 12-15 mice per genotype both gender, two-tailed Student’s t-test with Welsh correction *or # p < 0.05, ** p < 0.01, *** or ### p < 0.001, * comparison WT/WT to KI lines, # comparison WT/304Q to 304Q/304Q, black = WT/WT, dark green = WT/304Q, bright green = 304Q/304Q.

Gender separation of SC, PA, and BOS demonstrated similar tendencies in males and females, but in most cases did not reach significance potentially due to reduced animal numbers (Additional file 5: Figure S5).

### RNA-seq revealed differences in cerebellar gene expression starting at 2 months of age

RNA-seq of cerebellar samples of pre-symptomatic (2 months) and symptomatic (12 months) 304Q/304Q mice and their age-matched WT/WT littermates showed differential expression of ten genes at the pre-symptomatic time point and 398 differentially expressed genes (DEGs) at the symptomatic time point. Along the age dimension, 234 genes were differentially expressed comparing 2- and 12-month old WT/WT mice, and 334 genes comparing 2- and 12-month old 304Q/304Q mice (Figure 5A). Overlapping DEGs for pre-symptomatic and symptomatic mice, identified eight out of ten genes (*Car9, Ccdc63, Igfbp5, Il20rb, Il33, Syndig1l, Gm39465, 4930447F24Rik*) already affected at the pre-symptomatic stage, being still differentially regulated in the symptomatic stage, indicating an early onset in differential expression that is persistent over time (Figure 5B). For *Atxn3* itself we observed a downregulation in 2-month-old 304Q/304Q mice, while its gene expression stayed comparable to levels of 2-month-old 304Q/304Q mice in 12-month-old WT/WT and 304Q/304Q mice (Additional file 6: Figure S6A) and *Acy3* was upregulated only in 2-month-old 304Q/304Q mice, but not in 12-month-old ones. To further confirm changes in gene expression of *Acy3, Car9, Ccdc63, Igfbp5, Il20rb, Il33*, and *Syndig1l*, respectively, we performed qRT-PCRs for both time points in WT/WT, WT/304Q, and 304Q/304Q cerebellar samples. In 2-month-old mice (Additional file 6: Figure S6B) changes in gene expression did not reach significances using qRT-PCR. In the 12-month-old mice (Figure 5C), we confirmed the downregulation of *Car9, Ccdc63, Igfbp5*, and *Il33* as well as the upregulation of *Acy3* in 304Q/304Q cerebellar samples compared to WT/WT littermates. For all four downregulated genes, the heterozygous WT/304Q samples showed an intermediate level between those of WT/WT and 304Q/304Q. We tried to validate Il*20rb* and *Syndig1l* gene expression with several qRT primer pairs, but could not verify the results from the RNA-seq (data not shown). On protein level, we observed a significant increase in Acy3 at 12 months of age for WT/304Q and 304Q/304Q mice compared to WT/WT littermates, but no significant changes at 3 months of age (Figure 5D + 5E). For Car9 protein expression, we demonstrated that the amount of protein was significantly reduced or showed a tendency towards a reduction in both WT/304Q and 304Q/304Q mice compared to WT/WT mice at both time points (Figure 5F + 5G). Ccdc63 protein was significantly reduced at 3 months but not at 12 months of age (Figure 5H + 5I), and no differences were observed for the protein levels of Il33 (Additional file 6: Figure S6E + S6F). Igfbp5 protein was not detectable in 3- and 12-months-old mice independent from the genotype as well as in prenatal rat tissue used as control (data not shown). To specifically analyze full-length wildtype and expanded soluble Atxn3, highly-sensitive immunoassays, so-called TR-FRET assays, were used. Thereby, we observed an increase in total soluble Atxn3 in both, WT/304Q and 30Q/304Q mice, at 3 months of age, which was reduced to WT/WT level in 18-month-old mice (Additional file 6C). Expanded Atxn3 was increased at both time points in WT/304Q and 304Q/304Q mice compared to WT/WT littermates, but soluble protein levels were reduced at 18 months of age (Additional file 6: Figure S6D). Importantly, the strongest reduction was found in homozygous animals were the highest amounts of aggregates were detected (Figure 1B + 1C).

**Figure 5:**
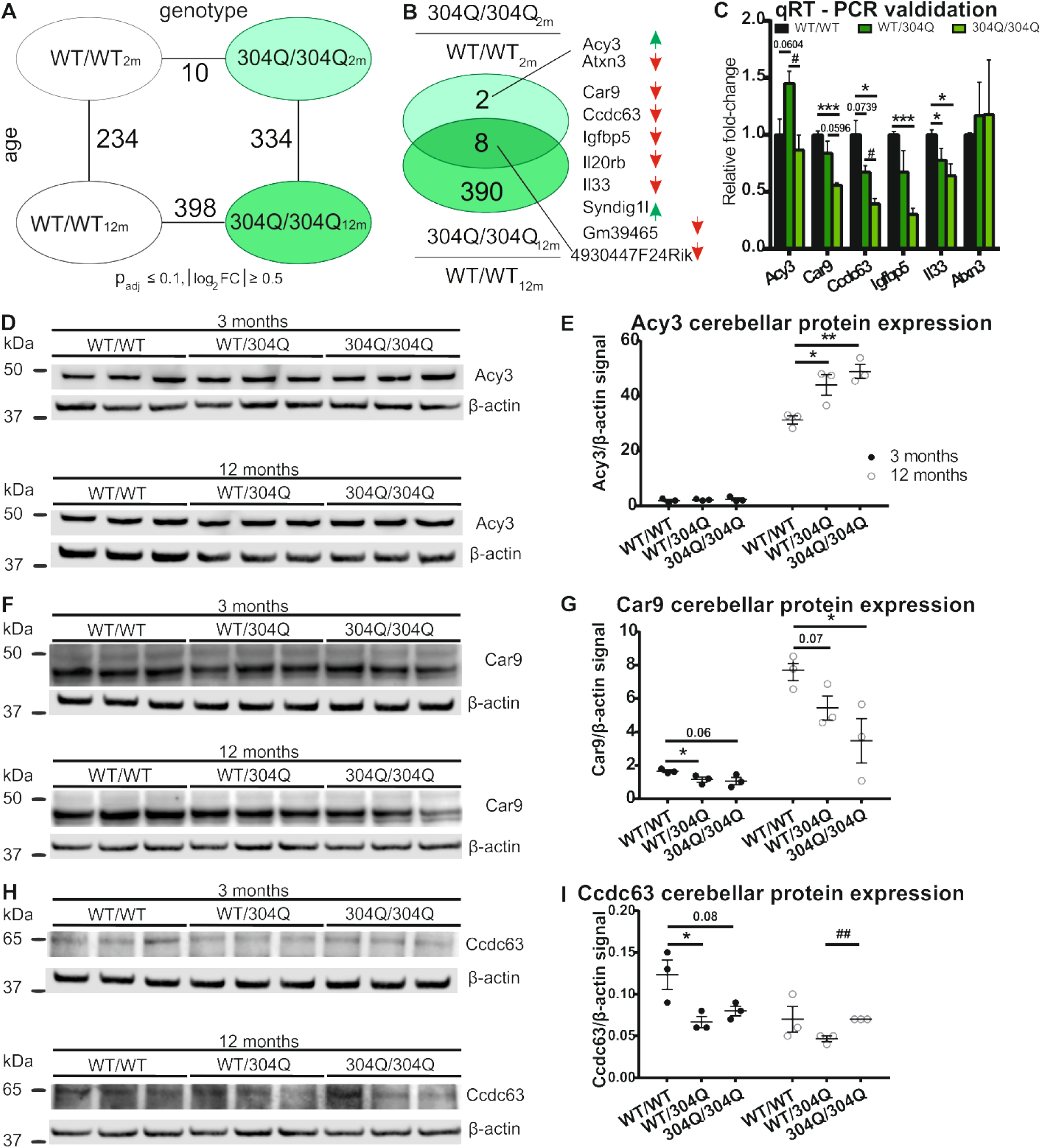
Altered gene and protein expression in SCA3 KI mice. (A) Total RNA from cerebellar tissue was isolated from 2- and 12-month-old WT/WT and 304Q/304Q mice for RNA-seq to interrogate transcriptomic changes. Primary contrasts along the genotype and age dimension revealed 10 DEGs in 2-month and 398 DEGs in 12-month-old animals. (B) Genotype effect revealed eight differentially expressed genes in 2- and 12-month-old 304Q/304Q KI mice. (C) qRT-PCR validation in RNA samples of 12-month-old mice confirmed significant downregulation of *Car9, Ccdc63, Igfbp5*, and *Il33* in 304Q/304Q and upregulation of *Acy3* in WT/304Q mice. No significant differences for *Atxn3* were detected. (D-E) Protein expression of Acy3 was unchanged with 3 months and elevated in KI mice with 12 months of age. (F-G) Total Car9 protein expression showed reduction of Car9 protein in both 3- and 12-month-old KI mice. (H-I) Ccdc63 protein expression was reduced in 3-month-old KI mice. In 12-month-old mice, Ccdc63 expression was reduced in WT/304Q compared to 304Q/304Q mice. RNA-seq n = 5; qRT-PCR/protein n = 3, two-tailed Student’s t-test * or # p < 0.05, ** or ## p < 0.01, * comparison WT/WT to WT/304Q and 304Q/304Q, # comparison WT/304Q to 304Q/304Q, 2m/12m = 2/12 months old animals. β-actin was used as loading control.

### Age-dependent gene expression changes arise from two distinctive perturbation modes

To better understand the age-dependent increase of expression perturbations in the cerebellum (Figure 5A) of the 304Q/304Q model, all 400 DEGs (union of 2- and 12-month-old contrasts) were visualized across the experimental groups (Figure 6A). Upon hierarchical clustering, these expression signatures partitioned into two primary classes. Class I comprises clusters 1 and 4, and class II comprises clusters 2 and 3 (Figure 6B). Genes in class I showed an age-dependent aggravation of expression perturbations in 304Q/304Q animals that did not occur in WT animals. In class II, the underlying genes underwent age-depended expression adaptations in WT animals that, however, failed in KI mice or, for some genes, were even oppositely regulated. Similar modes of age-dependent gene expression changes have already been described in a BAC SNCA mouse model for Parkinson’s disease and might represent a common pattern for neurodegenerative diseases (Hentrich *et al*. 2018).

**Figure 6:**
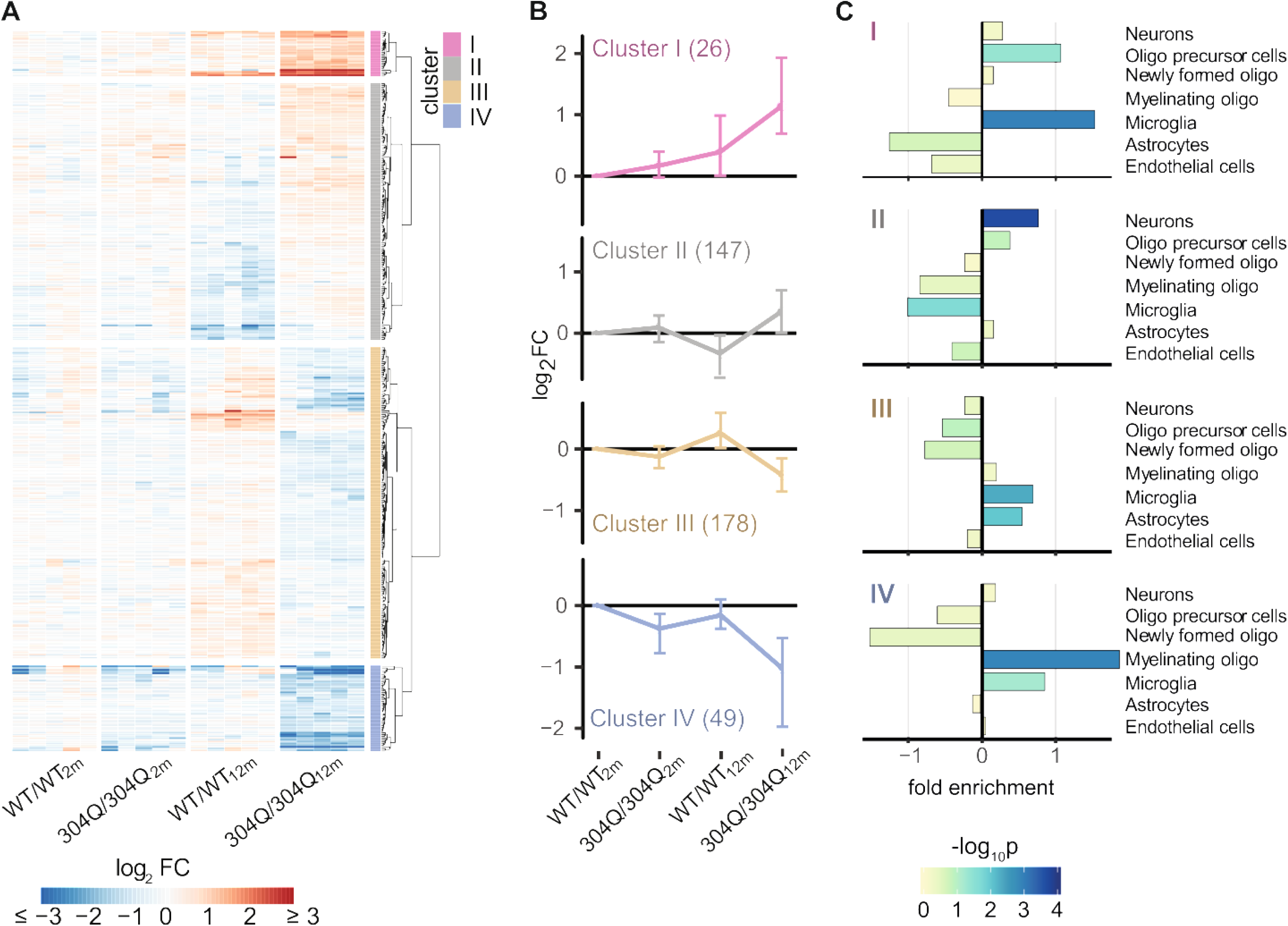
Modalities of age- and genotype-dependent gene expression perturbations. (A) Heatmap of hierarchically clustered expression levels (log_2_ expression change relative to WT/WT_2m_) across all experimental groups. Analyzed were the 400 DEGs found to be differentially expressed either in 2- or 12-month-old 304Q/304Q mice compared to their age-matched WT/WT littermates (see Figure 5B for comparison). Based on their gene expression pattern, the 400 DEGs partition into four main clusters. (B) Subplots of each gene cluster shown as expression centroid (±SD). (C) Enrichment of DEGs for cerebellar cell types for each cluster based on cell type-specific reference data (Zhang *et al*. 2014). Bars show fold enrichments with color-coded *p*-values of two-sided Fisher’s exact tests.

Class I was enriched for *structural constituent of myelin sheath* among Gene Ontology terms based on genes like *Mal, Plp1*, and *Mobp*. Class II was enriched for *voltage-gated cation channel activity* and comprises genes like *Kcnj3, Itar, Rest, Kcnk2, Cacna2d1*, and *Kcng4*.

The results for class I suggested a potential role of oligodendrocytes in the observed perturbations. To test this hypothesis, cell type-specific expression data from the Brain-Seq database (Zhang et al., 2014), which reports the cerebellar expression level of genes across seven cell types, were used to determine enrichments among the 400 DEGs. Indeed, class I, in particular the subset of DEGs in cluster 4, was enriched for genes attributed to myelinating oligodendrocytes (Figure 6C).

### Shared perturbances in oligodendrocytes between 304Q/304Q mice and SCA3 patients

In order to relate gene expression changes observed in mouse cerebellum to human, we interrogated *post-mortem* cerebellar tissue of SCA3 patients and healthy controls using RNA-seq. Of 2108 DEGs identified in humans, 69 were also differentially regulated in the KI mouse model (Figure 7A). These common DEGs were most significantly enriched for *structural constituent of myelin sheath* among molecular functions of Gene Ontology terms (Figure 7B). To further investigate this re-occurring result for a potential role of oligodendrocytes, the 69 DEGs were classified by their cell-type-specific expression according to the Brain-Seq database (Zhang et al., 2014). The strongest enrichment for 67 attributable genes was found for myelinating oligodendrocytes (Figure 7C). Seven out of ten genes underlying this term showed the same perturbation directionality when comparing 304Q/304Q KI mouse and SCA3 patient expression perturbations (Figure 7D).

**Figure 7:**
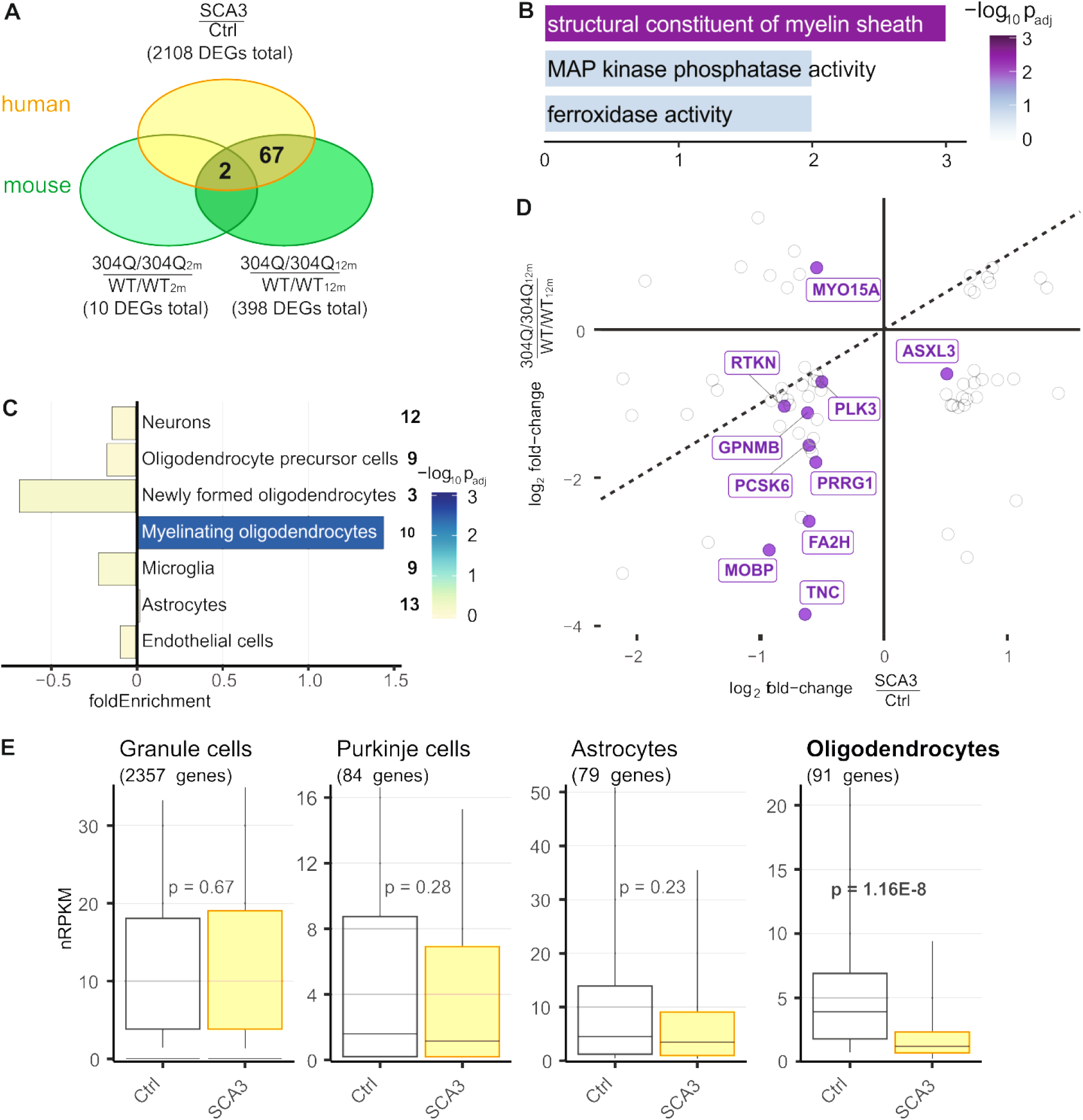
Common expression disturbances in KI mouse model and SCA3 patients point towards myelinating oligodendrocytes. (A) Venn diagram comparing DEGs in 2- and 12-month-old 304Q/304Q mice with orthologue DEGs found in the cerebellum of SCA3 patients. 69 genes were found to be differentially expressed in both mouse and human *post-mortem* cerebellum samples. (B) Fold enrichment of Gene Ontology terms (molecular function) for 69 common DEGs shared between KI mice and human patients. (C) Enrichment of shared DEGs for cerebellar cell types according to cell type-specific reference data (Ref Brain-Seq (Zhang *et al*. 2014)). Number of attributed genes indicated on the right. Bars show fold enrichments with color-coded *p*-values of two-sided Fisher’s exact tests. (D) Scatter plot of expression changes for 69 common DEGs identified in cerebellum of mouse and human. DEGs assigned to myelinating oligodendrocytes in purple. (E) Cell type-specific cerebellar gene expression in SCA3 patients and controls. Boxplots show geometric mean and 10th, 25th, 75th, and 90th quantile of nRPKM values for all genes attributed to distinct types based on human reference data (Kuhn *et al*. 2012). Number of genes per cell type in brackets. P-values of two-tailed Mann-Whitney U tests indicated for each cell type.

The common downregulation of genes (3^rd^ quadrant) attributed to oligodendrocytes prompted us to assess cell type composition effects. Using Brain-seq reference data first, the average expression of all genes attributed to myelinating oligodendrocytes indicated indeed a slightly reduced expression in 12-month-old 304Q/304Q mice (Additional file 7: Figure S7). Intriguingly, this reduction in oligodendrocytes was significant in SCA3 patients according to cell population-specific reference data for human cerebellum (Kuhn *et al*. 2012).

Taken together, these findings are in line with previous reports on KI/SCA3 models, which suggests an important role of oligodendrocytes (Ramani *et al*. 2017). Here, we now related these findings to humans and present evidence for impairment and reduction of oligodendrocytes in the cerebellum of SCA3 patients.

## Discussion

In this study, we generated and characterized a novel Atxn3 KI mouse model that depicts the SCA3 clinical and neuropathological phenotype in patients in an unprecedented way. With this new model, we could mimic neuropathological features, like the formation of aggregates and cell loss of Purkinje cells, but also reproduce a behavioral phenotype manifested by a reduction in body weight as well as balance and gait instability. Furthermore, transcriptional changes in this model revealed disturbances in the myelin sheaths and in myelinating oligodendrocytes, changes that were shared with perturbations in the cerebellum of *post-mortem* SCA3 patients (Figure 8). Therefore, this model is most suitable to investigate early-onset and longitudinal changes, making it an optimal candidate to study disease initiation, progression, and treatment.

**Figure 8:**
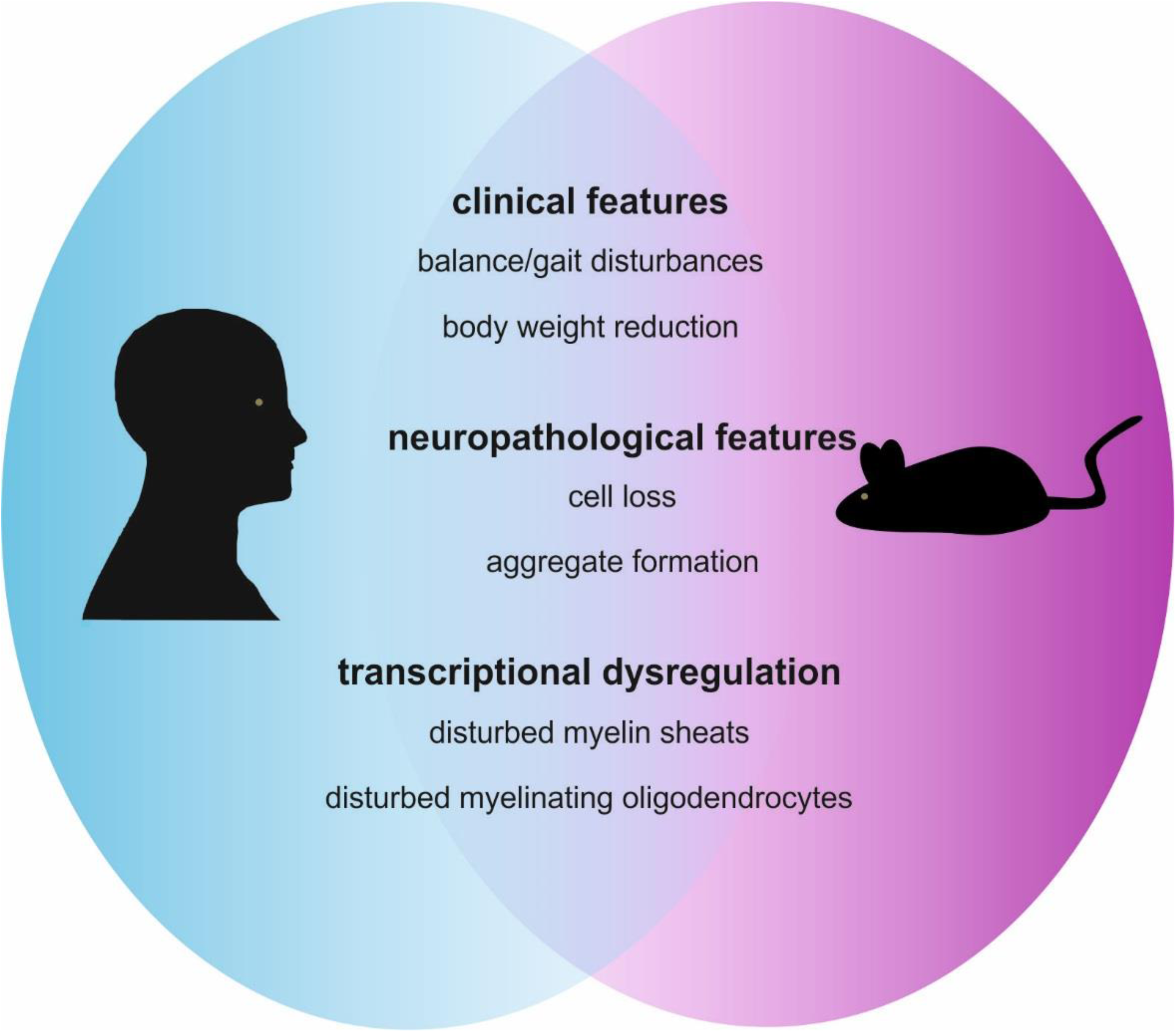
Summary of the common findings of SCA3 features in human patients and our Atxn3 KI mice. This novel Atxn3 KI mice share clinical features (balance and gait disturbances and reduced body weight), neuropathological features (cell loss and aggregate formation) and transcriptional dysregulation (disturbed myelin sheaths and myelinating oligodendrocytes) with human SCA3 patients.

It is known that KI models for SCA3 containing a CAG expansion within the range of repeat length expansions of human patients, display only a mild or no behavioral phenotype (Switonski *et al*. 2015, Ramani *et al*. 2017). The same is true for other polyQ diseases like SCA1 (Lorenzetti *et al*. 2000), SCA2 (Damrath *et al*. 2012), or HD (Menalled *et al*. 2002). Especially for the latter it has been proven, that a hyper-expansion of the repeat is necessary to trigger a behavioral phenotype (Wheeler *et al*. 2000). Moreover, the late-onset behavior phenotype in the hemizygous SCA3 YAC84Q model, which expresses a full-length copy of human cDNA containing 84 CAG repeats in the *ATXN3* gene with all regulatory elements (Cemal *et al*. 2002), was often not reproducible in follow-up studies (Toonen *et al*. 2018, Weber *et al*. 2020). This leads us to the assumption, that to trigger a phenotype comparable to human patients, a hyper-expansion of the polyQ tract is necessary. Therefore, we decided to generate a new SCA3 knock-in mouse model with a hyper-expansion of more than 150 glutamines. Further, we used an interrupted repeat, meaning a CAA instead of a CAG codon on every third position, both translated to glutamine on protein level. This was meant to increase the stability of the resulting polyQ tract (Menon *et al*. 2013), although we are aware that our mice could miss the feature of potential RNA toxicity, that is less prominent in interrupted repeats and can also contribute to disease progression (Li *et al*. 2008, Jung *et al*. 2011).

Protein expression analysis of our KI line revealed soluble Atxn3 protein expression throughout aging in the brain and several peripheral organs, including heart, lung, liver, and spleen, while we could not detect soluble Atxn3 in kidneys and muscle tissue. These findings are consistent with the protein expression pattern found by Switonski *et al*. (2015) in their KI line. Further, we observed a significant increase of Atxn3- and ubiquitin-positive aggregates in both WT/304Q and 304Q/304Q KI mice over time. This aggregate formation occurs already in early stages (first measured with 3 months of age), long before any behavioral changes get manifested. Moreover, we observed that more soluble Atxn3 was expressed in the KI mice with 3 months of age, but that level was reduced in 18 months old mice, indicating its inclusion into aggregates. The localization of these aggregates is ubiquitous in the brain, but especially prominent in the DCN, pons, hippocampus, and the cerebellar loops. All these areas are known to be vulnerable in SCA3 patients (Schmidt *et al*. 1998, Yamada *et al*. 2004). The localization of the aggregates is comparable to the findings of Ramani and colleagues (Ramani *et al*. 2015, Ramani *et al*. 2017) in their dupKi mouse. Another neuropathological variation we observed was a reduction and shrinking of Purkinje cells. Here, we could already detect a reduction in cell number and size in 3-month-old 304Q/304Q KI mice persistent with aging. Changes in Purkinje cell pattern and atrophy have been reported before, both for SCA3 mouse models (Hubener *et al*. 2013, Switonski *et al*. 2015) and also in histopathological examinations of SCA3 patients (Scherzed *et al*. 2012). Whether or not there is a connection between neuronal cell loss, cellular dysregulation, cerebellar function, and a motor phenotype is not completely clear yet. While Shakkottai *et al*. (2011) reported of changes in Purkinje cell firing concurring with behavioral deficits in YAC84Q mice even before observable neurodegeneration was detected, Costa Mdo *et al*. (2013) described behavioral deficits in the same mouse model, but could not detect changes in Purkinje cell count. Toonen *et al*. (2018) did not observe behavioral differences at the same time point. Therefore, the relation between Purkinje cell loss and a motor phenotype needs further investigation in several SCA3 models.

These neuropathological features described above also have an impact on the phenotypical development of the KI mice. In male mice, we observed early reduction of body weight in heterozygous and homozygous animals, whereas the WT/304Q mice became later significantly different compared to their WT/WT littermates and were not as severely affected as the 304Q/304Q mice. Female WT/304Q KI mice were not affected by body weight reduction, and 304Q/304Q females were significantly reduced compared to WT/WT littermates, but not nearly as strong as the males. Body weight reduction has been shown for the hemizygous YAC84Q mice (Cemal *et al*. 2002) and is consistent with SCA3 patients, where a negative correlation between CAG repeat length and body weight has been proven (Saute *et al*. 2012, Saute *et al*. 2012). The discrepancy we noticed between male and female mice has also been reported for KI and transgenic mice of other polyQ diseases. In Huntington’s research, for example, male transgenic mice and rats, as well as the CAG140 KI mice, are more severely affected by body weight reduction than their female littermates (Bode *et al*. 2008, Phan *et al*. 2009). These differences in body weight affect also later experiments. Therefore, we observed physiological changes like size reduction and hunchback formation mainly in male mice. During RotaRod experiments, homozygous male mice seem to benefit from the reduced body weight at intermediate time points and become comparable to WT/WT littermates in later time points. However, in female mice, which are less affected by changes in body weight, we observed a tendency to perform worse in this motor task from the beginning and being significantly worse at late stages. These gender discrepancies have to be kept in mind for further studies. However, we do not recommend working with male or female mice only, when dealing with a disease in which male and female patients are impaired equally. Other behavioral experimental setups, like the Catwalk experiment, were seemingly unhindered by body weight changes and therefore we did not observe significant gender differences. Here, we detected increased step cycles in front and hind paws, decreased print area in front and hind paws, and increased base of support for the hind paws. Results were always significant in 304Q/304Q mice and for the base of support and the step cycle in the hind paws also for WT/304Q mice. These findings are comparable to the ataxic phenotype in human patients, where differences in posturing for better balance are also described (Riess *et al*. 2008).

These measurable and quantifiable parameters, like aggregate formation, weight reduction, and motor phenotype, render these KI mice as an optimal model system for therapy studies or biomarker validations. First studies have already used this KI model to prove antisense oligonucleotide (ASOs) therapy and longitudinal biomarker studies. Martier *et al*. (2019) could recently show that they can reduce the amount of expanded *Atxn3* mRNA in these KI mice by delivering microRNAs in an adeno-associated-vector system. Moreover, Wilke and colleagues are using this model to investigate the progression of neurofilament light chain (NfL) and phosphorylated neurofilament heavy chain (pNfH) as potential easily accessible serum biomarkers in a longitudinal study with mice in the age range from 2 to 24 months. They could prove the increase of the biomarkers already in 6-month-old mice, long before any symptoms developed and were able to correlate their findings with the biomarker levels in human patients (Wilke *et al*. 2020, medRxiv, doi: https://doi.org/10.1101/19011882). Additionally, it was possible to track first abnormalities in walking behavior by DeepLabCut software based on deep learning networks, already in 9-month-old homozygous KI mice, a time point before motor deficits were detectable with other behavioral experiments (Lang *et al*, 2020; submitted as conference manuscript to the IEEE Engineering in Medicine and Biology Society). These findings support the great potential of this KI mouse line for further biomarker or treatment studies.

Potential biomarkers could also arise from high powered transcriptomic studies. In line with two other SCA3 rodent models, *Interleukin-33 (Il33)* is persistently downregulated in cerebellar samples of the 304Q/304Q KI mice at both investigated time points (2 and 12-months of age). In YAC84Q mice the *Il33* transcript is also downregulated in brain stem samples of 6-month-old SCA3 mice (Ramani *et al*. 2017) and whole brain samples of 17.5-month-old mice (Toonen *et al*. 2018). In addition to this, *Il33* shows highest expression levels in the central nervous system, being constitutively expressed in astrocytes, microglia, and oligodendrocytes (Liew *et al*. 2016). Moreover, a deficiency for *Il33* is associated with neurodegeneration and impaired repair of neurons in aged mice accompanied by memory loss and cognitive impairment, resembling an Alzheimer’s disease-like phenotype, including tau abnormality (Carlock *et al*. 2017).

Another candidate gene as a potential transcriptional biomarker is *Aminoacyclase-3 (Acy3)*. It was upregulated not only in our 2-month-old KI mice, but also in 6-month-old YAC84Q mice line (Ramani *et al*. 2017), where they showed an upregulation of protein expression in brain samples of human SCA3 patients compared to Alzheimer’s patients as control. Further, this gene has been shown to be robustly upregulated in several transcriptomic studies on HD (Becanovic *et al*. 2010, Giles *et al*. 2012, Langfelder *et al*. 2016). The most interesting aspect about Acy3 as biomarker lies in the fact that this enzyme breaks down N-acetyl aspartate (NAA) a commonly used marker in magnetic resonance imaging, which is decreased not only in HD but also several SCAs (Sanchez-Pernaute *et al*. 1999, Lopes *et al*. 2013, Klaes *et al*. 2016). Therefore, also NAA is worthy to be investigated as a useful imaging biomarker for SCA3.

Not only are Acy3 and Il33 associated with oligodendrocytes, but also our cell-type-specific data analysis point at disturbances in myelinating oligodendrocytes and myelinating sheaths. Importantly, these gene expression changes were shared between our SCA3 KI mice and human *post-mortem* cerebellar SCA3 patient tissue. A high number of downregulated genes in both, the KI mice and humans were enriched for this cell type. These findings are in line with the transcriptomic studies of Ramani *et al*. (2017) and Toonen *et al*. (2018) in the YAC84Q mice, with oligodendrocytes appearing mainly affected by transcriptional changes. Ramani and colleagues further showed that ATXN3 is present in the cell nuclei of oligodendrocytes. In a later study McLoughlin et al. (2018) demonstrated, that these oligodendrocytes can be directly targeted by antisense-oligonucleotides, reversing changes in gene expression, e.g. the upregulation of *Acy3* and other oligodendrocyte specific genes. In SCA3 patients, the aggregation of ATXN3 in oligodendrocytes has yet not been investigated, but ATXN3 aggregation in Schwann cells, the myelinating cells of the peripheral nervous system, of SCA3 patients has been proven (Suga *et al*. 2014). White matter changes have also been reported in other polyQ diseases. Jin *et al*. (2015) were able to show early myelination defects in a Q250 mouse model of HD, possibly caused by altered oligodendrocyte differentiation. Further, the overexpression of huntingtin in oligodendrocytes provokes behavioral deficits and increased demyelination in mice (Huang *et al*. 2015). Up until now, white matter changes have been assumed to arise secondary after neuronal loss occurred. However, this study together with others supports the hypothesis that the expression of expanded Atxn3 in oligodendrocytes might impair proper myelination in parallel to or even proceeding neuronal cell damage.

## Conclusion

Taken together, we present a novel SCA3 Atxn3 KI mouse model that represents the human patient phenotype on a neuropathological, behavioral, and transcriptomic level and is, therefore, an ideal system for further studies, including therapy and biomarker investigations.

## Acknowledgment

We would like to thank Dr. Jonasz Weber, Dr. Jana Schmidt, Dr. Libo Yu-Täger and Alexandra Grenzendorf for their advice and knowledge as well as providing protein samples for establishing of certain techniques and Dr. Manuela Neumann and Johannes Hanselmann of the DZNE Tuebingen, for assisting with the overview images of the mice brains. Further, we want to thank Wilfred F.A. den Dunnen, Department of Pathology and Medical Biology, University Medical Center Groningen, University of Groningen, Groningen, The Netherlands for his contribution in collecting human brain tissue samples.

## Funding

The research leading to these results has received funding from the European Community’s Seventh Framework Program (FP7/2007-2013) under grant agreement 2012-305121 “Integrated European-omics research project for diagnosis and therapy in rare neuromuscular and neurodegenerative diseases (NEUROMICS).” Further support was received from JPND joint program for neurodegenerative disease, funding the project “Identification of genes that modulate the severity of neurodegenerative diseases (NeuroGem)” (FKZ01ED1507), the Netherlands Organization for Health Research and Development (ZonMw; AE, JM) and was supported by the Deutsche Forschungsgemeinschaft, DFG through the funding of the NGS Competence Center Tübingen (NCCT-DFG, project 407494995).

## Contribution to the study

**E.H**.: design and conceptualization of the study, acquisition of data, analysis of the data, drafting and, revision of the manuscript

**R.I**.: conduction of animal experiments and acquisition of data, analysis of the data, revision of the manuscript

**T.H**.: analysis of RNA sequencing data, revision of the manuscript

**Y.M**.: conduction of animal experiments and acquisition of data, analysis of the data, revision of the manuscript

**T.S**.: knock-in vector design, revision of the manuscript

**F.Z**.: generation of knock-in founder mice, revision of the manuscript

**N.C**.: conducting RNA sequencing experiments, revision of the manuscript

**E.A**.: acquisition of human brain material for protein and RNA studies, revision of the manuscript

**O.R**.: design and conceptualization of the study, revision of the manuscript

**J.S.-H**.: analysis of RNA sequencing data, drafting, and revision of the manuscript

**J.H-S**.: first experimental designs, design and conceptualization of the study, revision of the manuscript

---

**Figure S1:**
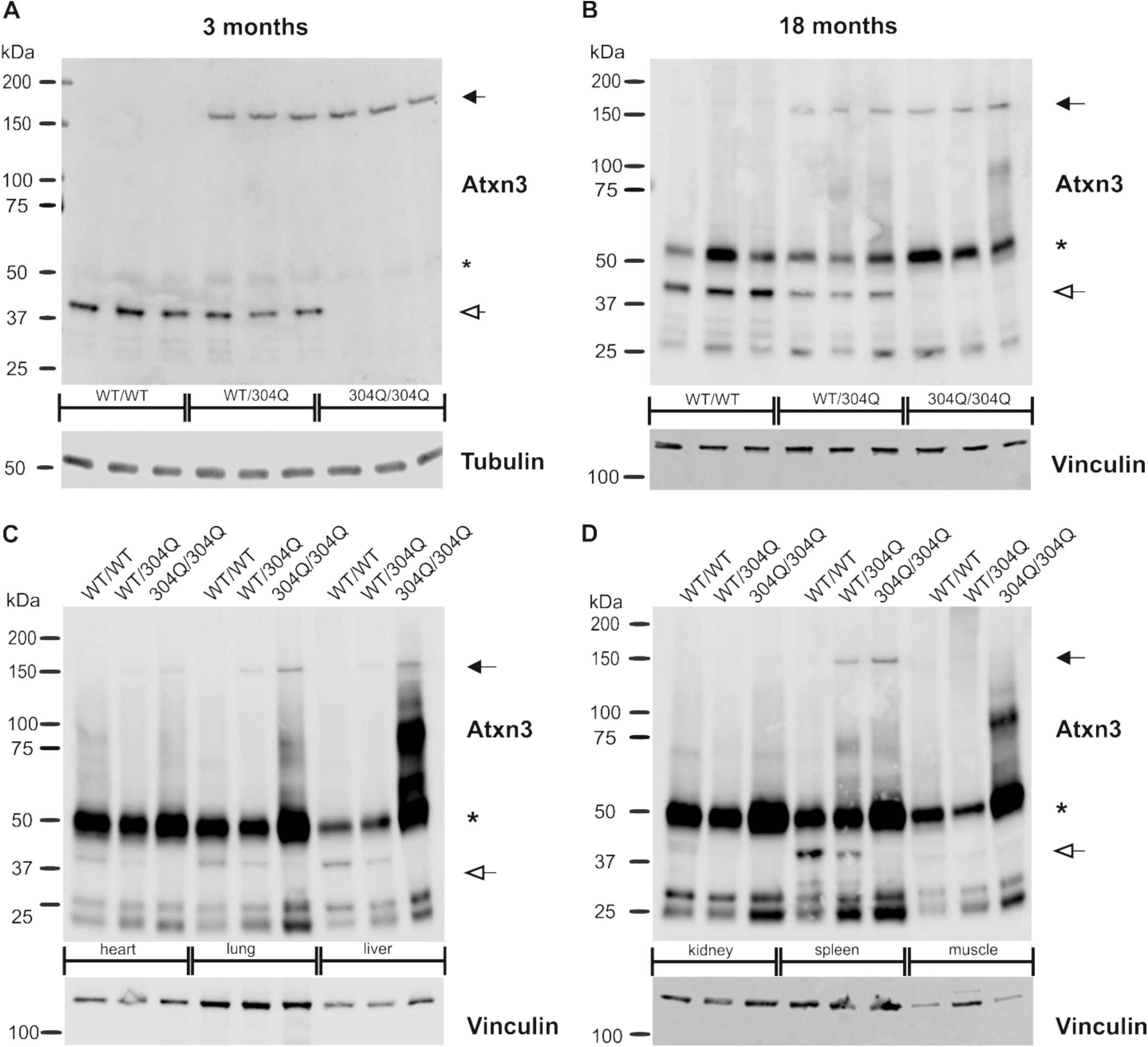
Protein expression of expanded Atxn3 in 3- and 18-month-old mouse brains and peripheral organs. (A-B) Western Blot analyses of whole brain lysates showed the expression of the polyQ-expanded and non-expanded Atxn3 in the heterozygous and homozygous KI lines WT/304Q and 304Q/304Q with 3 months (A) and 18 months of age (B). (C-D) Expression of non-expanded and polyQ-expanded Atxn3 in KI line 304Q is detectable in heart, lung, liver (C) and spleen (D), in muscle and kidney tissue no detection of expanded Atxn3 is possible (D). (A-B) n = 3 per genotype, (C-D) n = 1 per genotype; filled arrow = polyQ-expanded Atxn3, unfilled arrow = non-expanded Atxn3, * immunoglobulin protein detection. Vinculin or β-tubulin were used as loading control.

**Figure S2:**
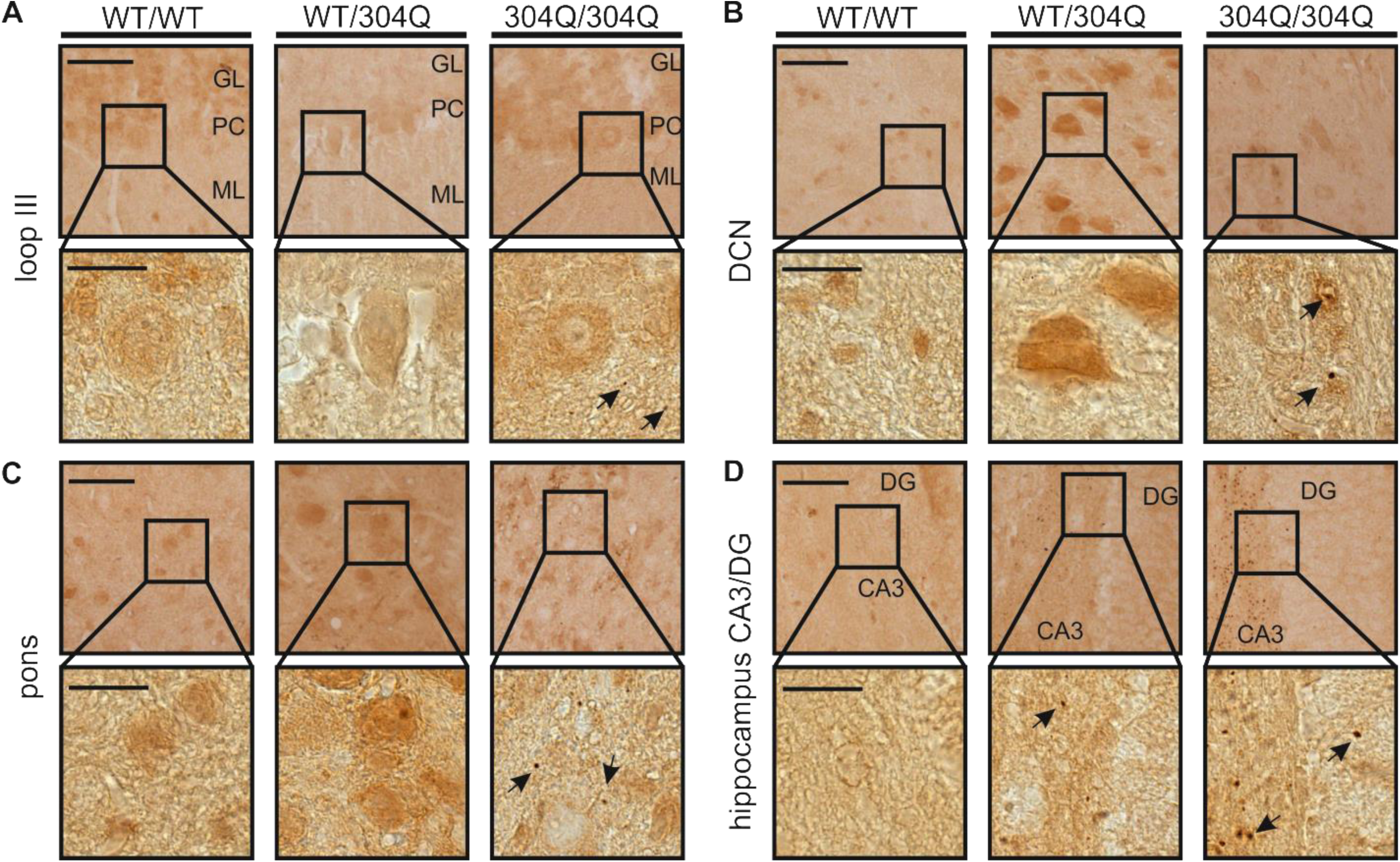
Beginning of aggregation of expanded Atxn3 in 3-month-old SCA3 KI mouse brain. Immunohistochemical (IHC) staining using Atxn3-specific antibody (clone 1H9) showed aggregate formation in WT/304Q and 304Q/304Q mice. IHC of 3-month-old mouse brains revealed increased diffuse nuclear staining in WT/304Q in the molecular layer of loop III (A), the DCN (B) and the pons (C) and beginning of aggregate formation in the hippocampus (D). In 304Q/304Q mice first aggregates in the molecular layer of loop III, DCN, pons, and hippocampus (A-D) can be observed at 3 months of age. Scale bar = 50 μm, inset scale bar = 20 μm. Atxn3 positive aggregates are indicated by arrows. n=3; GL = granular layer, PC = Purkinje cells, ML = molecular layer; DG = dentate gyrus, DCN = deep cerebellar nuclei..

**Figure S3:**
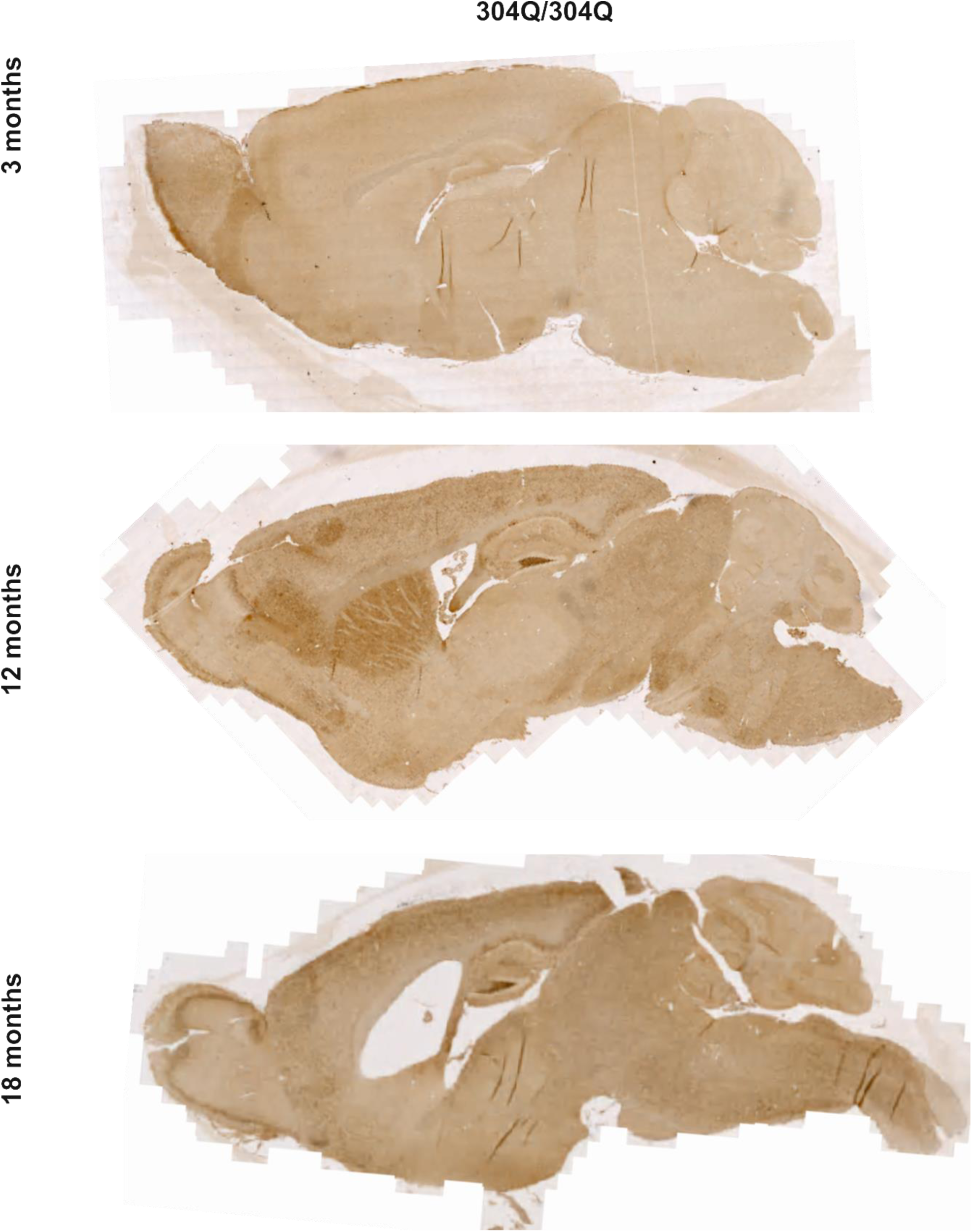
Overview of 304Q/304Q sagittal sections with 3, 12 and 18 months of age. Aggregate formation increased over time in 304Q/304Q mice. The increase is detectable especially in the olfactory bulb, cerebral cortex, hippocampus, cerebellum, and hindbrain by darkening of the tissue as the number of Atxn3 puncta increases.

**Figure S4:**
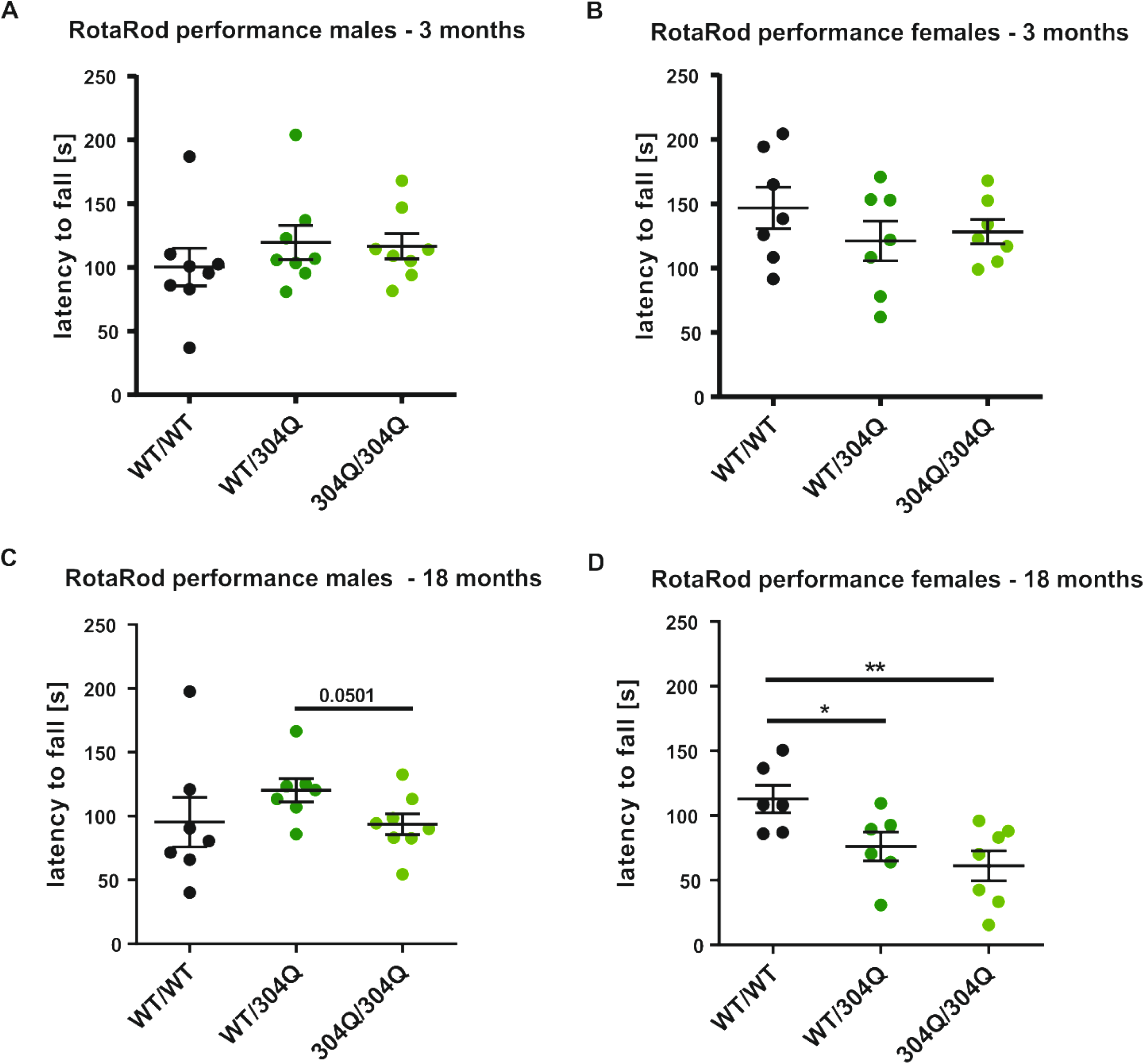
Coordination on RotaRod in male and female SCA3 KI mice. At 3 months of age, no significant differences in RotaRod performance were observed in either male (A) or female (B) SCA3 KI mice. (C) At 18 months of age, a tendency in homozygous 304Q/304Q male mice was observed to perform worse than their heterozygous littermates. (D) At 18 months of age, in WT/304Q and 304Q/304Q female mice coordination was significantly decreased compared to WT/WT littermates. n = 6-8 mice per genotype and gender, two-tailed Student’s t-test with Welsh correction * p < 0.05, ## p < 0.01 *** p < 0.001, * comparison WT/WT to KI lines, # comparison WT/304Q to 304Q/304Q, black = WT/WT, dark green = WT/304Q, bright green = 304Q/304Q.

**Figure S5:**
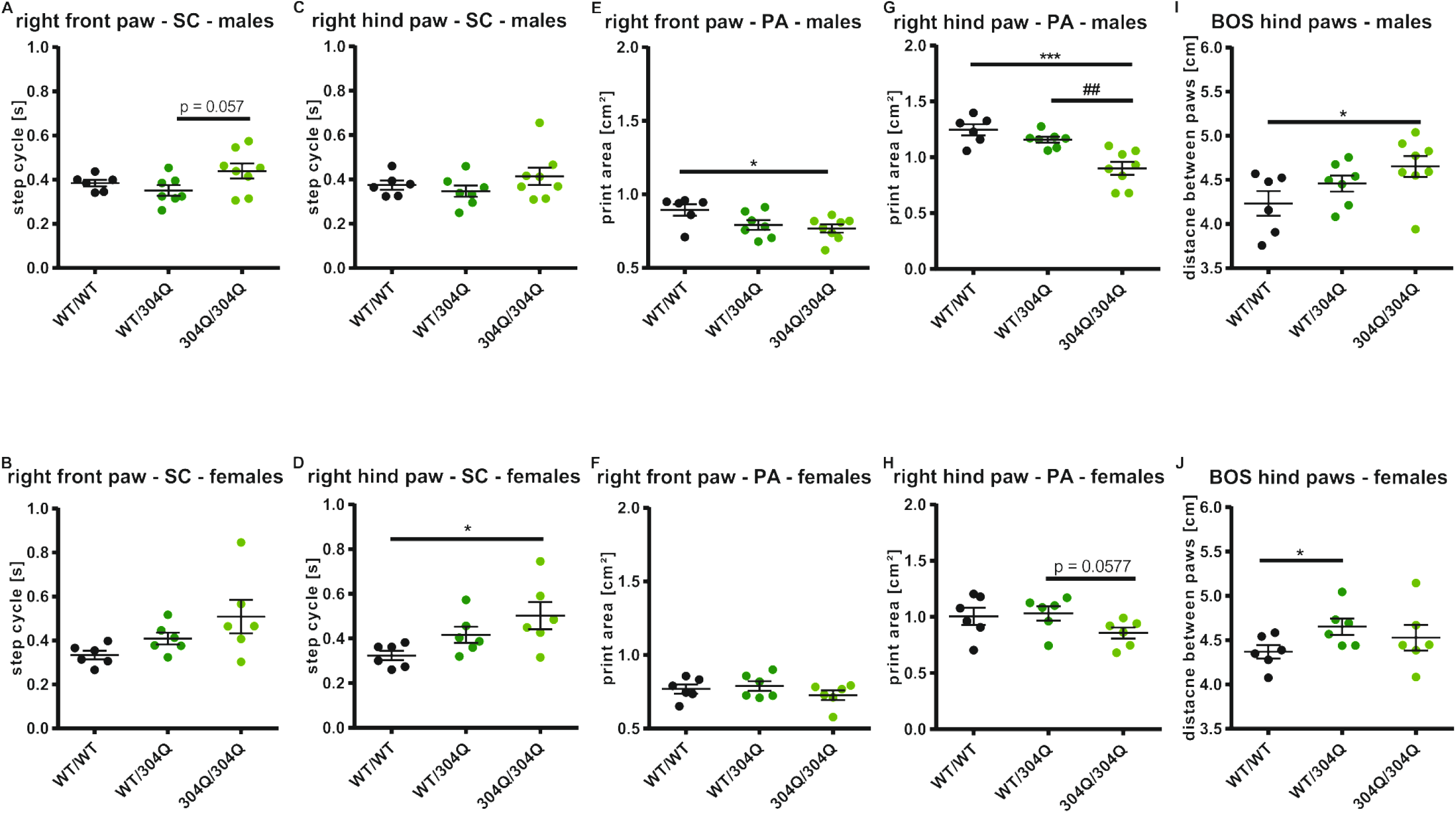
Altered paw positioning in old WT/304Q and 304Q/304Q KI mice separated by gender. (A-J) Gait analyses separated by gender revealed less significant impairment for WT/304Q and 304Q/304Q mice compared to their WT/WT littermates as in the pooled cohort. (A+C) In male mice step cycle (SC) was not increased in either right front (A) or right hind paw (B). For the front paw (A) a tendency for an increased SC was detectable between heterozygous and homozygous mice. (B+D) In female mice SC was not increased in the right front (B), but in the right hind paw (D) SC increased significantly in 304Q/304Q mice compared to WT/WT littermate. (E+G) Print area (PA) of the right front (E) and right hind paw (G) was significantly reduced in 304/304Q male mice compared to WT/WT littermates. For the hind paw this reduction was also significantly different to WT/304Q littermates. (F+H) In female mice no significant reduction in PA of the right front (F) and right hind paw (H) were detected. For the hind paw, a tendency for a reduction was observed in 304Q/304Q mice compared to their heterozygous littermates. (I-J) Base of support (BOS) of the hind paws was significantly increased in 304Q/304Q compared to WT/WT male littermates (I) and in WT/304Q female mice compared to WT/WT littermates (J). n = 6-8 mice per genotype and gender, two-tailed Student’s t-test with Welsh correction * p < 0.05, ## p < 0.01 *** p < 0.001, * comparison WT/WT to KI lines, # comparison WT/304Q to 304Q/304Q, black = WT/WT, dark green = WT/304Q, bright green = 304Q/304Q.

**Figure S6:**
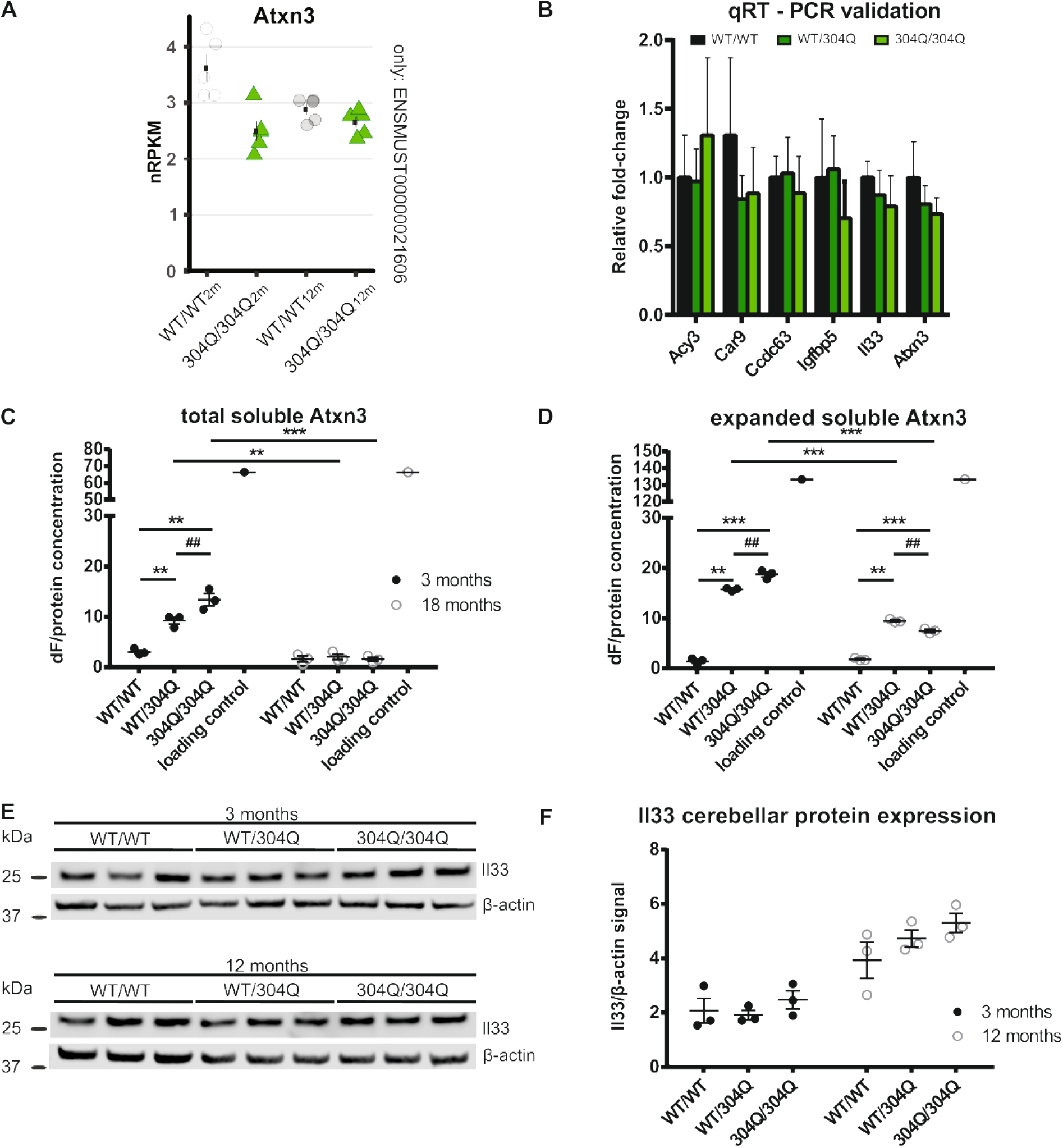
Altered gene and protein expression in SCA3 KI mice. (A) RNA-seq expression for Atxn3 showed increased gene expression in 2-month-old WT/WT mice and lower expression in all other groups. Expression is shown as normalized reads per kilobase per million total reads (nRPKM). (B) qRT-PCR validation in RNA samples of 2-month-old mice did not confirm significant down- and up-regulation of candidate genes. (C-D) Protein expression of total soluble Atxn3 (C) showed a significant increase in young WT/304Q and 304Q/304Q mice compared to WT/WT littermates. For the 304/304Q mice, the increase is also significant compared to WT/304Q mice. With 18 months of age, no differences between the genotypes were observed. But in WT/304Q and 304Q/304Q mice, the amount of total soluble Atxn3 was reduced compared to the same genotypes with 3 months of age. The amounts of expanded soluble Atxn3 were increased in WT/304Q mice to WT/WT littermates and in 304Q/304Q mice compared to WT/WT and WT/304Q mice in 3 and 18 months old mice. With 18 months of age, the amount of expanded soluble Atxn3 was reduced compared to the same genotypes with 3 months of age. (E-F) *Il33* gene expression is increased in 12-month-old compared to 3-month-old mice, but no genotype-dependent differences in the protein expression were found. RNA-seq n = 5; qRT-PCR and protein analysis n = 3, two-tailed Student’s t-test * or # p < 0.05, *** or ### p = 0.001, 2m = 2 months and 12m = 12 months old animals.

**Figure S7:**
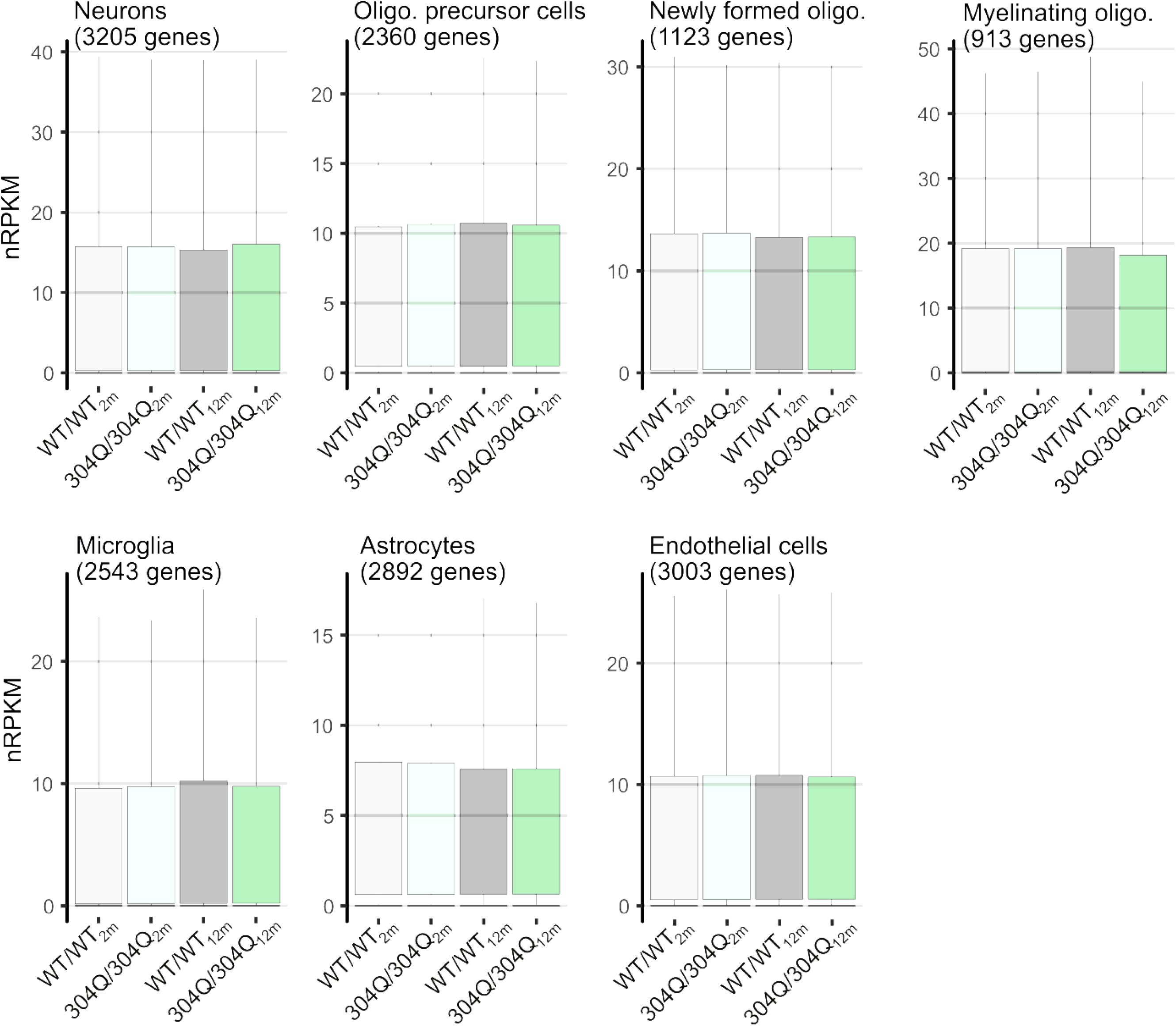
Cell type-specific cerebellar gene expression in 2- and 12-month-old WT/WT and 304Q/304Q animals. Boxplots show geometric mean as well as 10th, 25th, 75th, and 90th quantile of nRPKM values for all genes attributed to distinct cell types based on Brain-seq data (Zhang *et al*. 2014). Number of genes per cell type in brackets. No significant compositional changes were observed (Mann-Whitney U test, two-tailed).

